# STReTCh: a strategy for facile detection of mechanical forces across proteins in cells

**DOI:** 10.1101/2021.12.31.474658

**Authors:** Brian L. Zhong, Vipul T. Vachharajani, Alexander R. Dunn

## Abstract

Numerous proteins experience and respond to mechanical forces as an integral part of their cellular functions, but measuring these forces remains a practical challenge. Here, we present a compact, 11 kDa molecular tension sensor termed STReTCh (Sensing Tension by Reactive Tag Characterization). Unlike existing genetically encoded tension sensors, STReTCh does not rely on experimentally demanding Förster resonance energy transfer (FRET)-based measurements and is compatible with typical fix-and-stain protocols. Using a magnetic tweezers assay, we calibrate the STReTCh module and show that it responds to physiologically relevant, piconewton forces. As proof-of-concept, we use an extracellular STReTCh-based sensor to visualize cell-generated forces at integrin-based adhesion complexes. In addition, we incorporate STReTCh into vinculin, a cytoskeletal adaptor protein, and show that STReTCh reports on forces transmitted between the cytoskeleton and cellular adhesion complexes. These data illustrate the utility of STReTCh as a broadly applicable tool for the measurement molecular-scale forces in biological systems.

## INTRODUCTION

The ability of living cells to generate and sense molecular-scale mechanical forces is crucial to embryonic development, cell motility, and other biological processes. Nanoscale mechanical forces such as those produced by molecular motors typically range from single to tens of piconewtons (pN)^1-6^. Existing methods for characterizing forces experienced by proteins in intact cells rely on Förster Resonance Energy Transfer (FRET) or similar phenomena, where energy transfer between a pair of fluorophores flanking a molecular spring of known force constant is used to infer the extension of the spring and the force across the tension sensor^3, 7, 8^. While these sensors have yielded important biological insights^3-6, 8^, quantitative FRET measurements require specialized equipment, involved analysis techniques, and careful control measurements^9^, factors that limit the broad accessibility of such sensors. Consequently, existing molecular tension sensors are often used for qualitative assessments of relative forces. Moreover, FRET-based tension sensing modules are relatively large at ∼60 kDa in size, increasing the likelihood that their insertion may disrupt the function of the host protein. DNA hairpin-based tension sensors^10, 11^ represent another well-established modality for high-precision measurements of molecular mechanical forces, but such sensors are not currently viable for in-cell measurements.

Here, we present a 11 kDa molecular tension sensor module, which we term STReTCh (**S**ensing **T**ension by **Re**active **T**ag **Ch**aracterization), that is sensitive to forces in the single pN range and whose use follows typical fix-and-stain protocols and does not rely on FRET. We characterize the biophysical behavior of STReTCh under mechanical force and demonstrate the utility of STReTCh as a sensor of biologically relevant forces both at cell-substrate interfaces and inside cells.

## RESULTS

### Design and characterization of a novel force sensor

As a general approach, we sought to embed an epitope tag into a larger mechanosensitive domain such that the tag could not be recognized by its binding partner unless the domain was unfolded. We reasoned that the I10 immunoglobulin-like domain from human titin, which reversibly folds and unfolds at ∼6 pN on the seconds time scale^12^, could serve as the basis for a sensor of the pN-scale forces across individual proteins inside of cells. Specifically, we inserted the 13-amino acid SpyTag in the C-terminal loop of titin I10 to generate the STReTCh tension sensor (**Fig. 1**). SpyTag forms a covalent, isopeptide bond with its binding partner SpyCatcher^13, 14^. Thus, a fluorescently labeled SpyCatcher can be used as a labeling agent for the force-unfolded STReTCh sensor in a similar manner as antibodies for immunocytochemistry.

**Figure 1.**
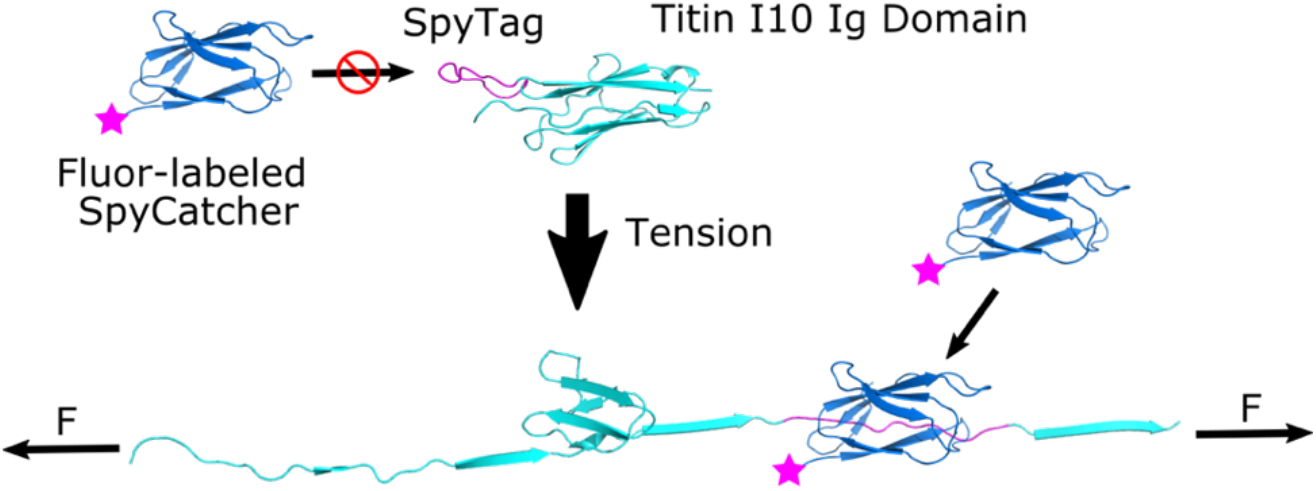
Design and operation of the STReTCh tension-sensing module. SpyTag was inserted into the Titin I10 domain such that SpyTag is only recognized by SpyCatcher when STReTCh is unfolded under tension. Unfolded STReTCh can be visualized using fluorophore-labeled SpyCatcher. The structure of STReTCh was predicted using RosettaRemodel based on the structure of Titin I10 (PDB: 4QEG).

Initial *in vitro* characterization of the STReTCh sensor revealed a lower melting temperature than that of the unmodified titin I10 domain (**Supp. Fig. 1**). To better characterize the unfolding behavior of STReTCh under tension, we used magnetic tweezers to apply tension in the single-pN range to individual surface-tethered STReTCh molecules (**Fig. 2a**). Individual STReTCh molecules exhibited reversible, steplike unfolding transitions of ∼10-15 nm (**Fig. 2b**) between folded and unfolded states, consistent with previous characterizations of titin I10 unfolding^12^. These transitions occurred at forces below 2 pN, consistent with the lower observed melting temperature relative to titin I10 (**Fig. 2b**). The fraction of time spent in the folded state for STReTCh molecules at various forces are plotted in **Figure 2c**, where the size of each data point corresponds to the total duration of time over which the molecule was observed. We fit the folded and unfolded state lifetimes measured using magnetic tweezers to a Bell-Evans kinetic model^15^ to obtain a maximum-likelihood estimate of the threshold unfolding force for STReTCh, which we found to be ∼1 pN. STReTCh is thus among the most sensitive molecular force sensors described to date, and capable of responding to the full range of biologically relevant, molecule-scale forces, which span 2 to ∼100 pN^3-6, 8, 10, 16, 17^.

**Figure 2.**
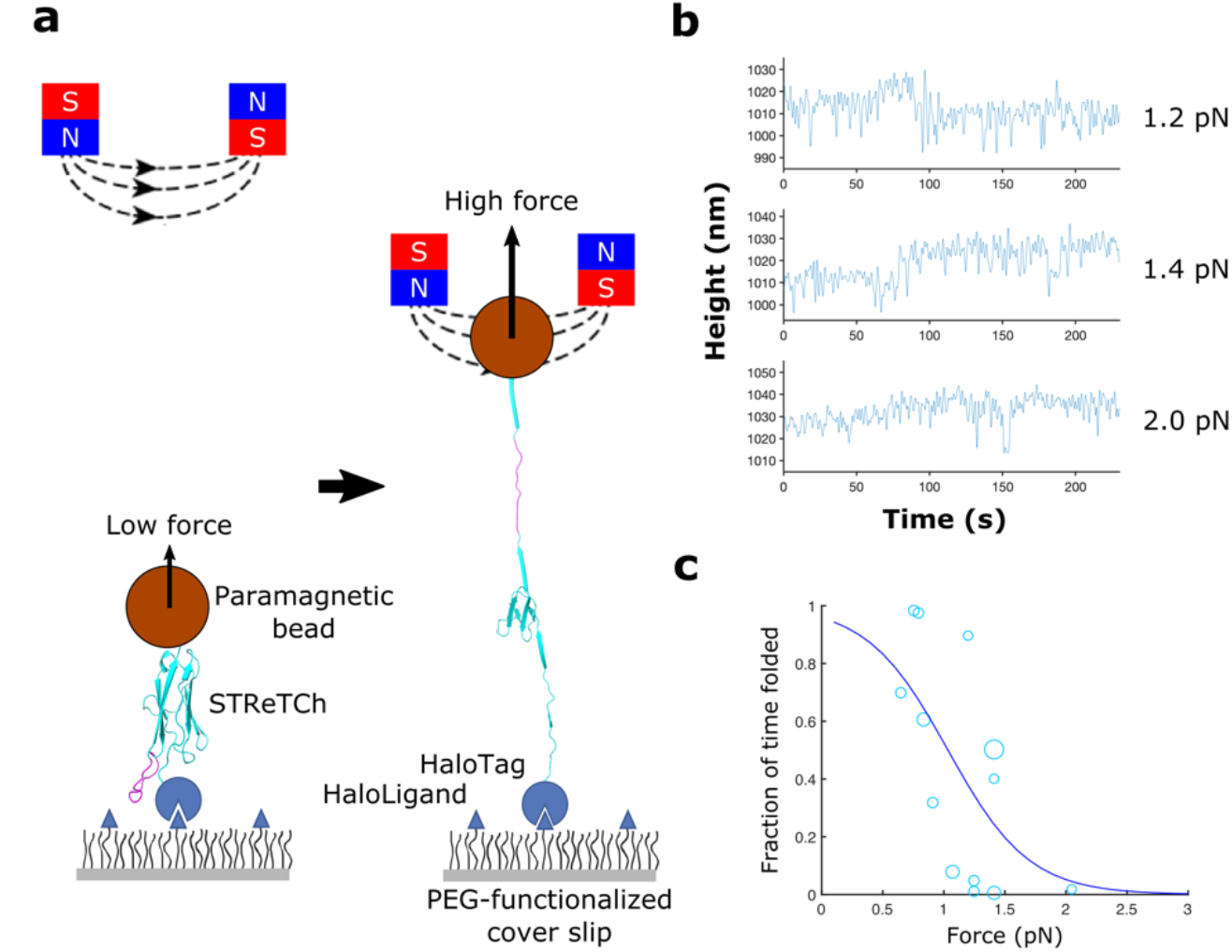
The STReTCh module is largely unfolded at forces above 1 pN. **(a)** Single molecule magnetic tweezers assay (not to scale). STReTCh molecules are covalently tethered to a microscope coverslip and bound at the other end to paramagnetic beads. Forces in the pN range are applied via a magnetic field gradient created by two permanent magnets, with increasing force as the magnets are brought closer to the sample (see Supp. Fig. 2b). **(b)** Sample traces of STReTCh under fixed levels of tension. STReTCh exhibits steplike changes in length of ∼10-15 nm at forces of 1-2 pN. Data shown were acquired at 50 Hz and filtered with a 7^th^ order Butterworth filter with a cutoff frequency of 0.5 Hz. **(c)** Unfolding behavior of STReTCh under mechanical force suggests that STReTCh transitions from primarily folded to unfolded at 1 pN. Each data point represents one trace acquired at the corresponding force, with the total dataset drawn from 7 distinct molecules measured across 6 independent experiments. The size of each marker is proportional to the total duration of the corresponding trace. The curve represents the Bell-Evans model fit to data. Parameters for the Bell-Evans model and corresponding confidence intervals are listed in Supp. Table 1.

### Measuring extracellular integrin-ECM forces using STReTCh

As a functional test of the STReTCh sensor, we evaluated whether STReTCh could detect forces generated at integrin-based cellular adhesions, hereafter generically termed focal adhesions (FAs), to the extracellular matrix (ECM)^18, 19^. We seeded HFFs transfected with focal adhesion marker GFP-paxillin (GFP-Pxn) onto a substrate functionalized with the STReTCh sensor fused to a C-terminal fibronectin-derived RGD peptide that mediates integrin-based adhesion^20^. We hypothesized that FA-localized forces exerted by HFFs would preferentially unfold STReTCh at FAs. After allowing the HFFs to adhere to the substrate, we treated the HFFs with 0.5% paraformaldehyde (PFA) to temporarily halt FA turnover^21^ prior to imaging and “stained” the sample with a truncated, minimal SpyCatcher^14^ labeled with Alexa Fluor 647 (647-SC) to visualize unfolded sensors. Following washout of 647-SC, the cells were imaged at 37 °C using total internal reflectance fluorescence (TIRF) microscopy (**Figure 3a, b**).

**Figure 3.**
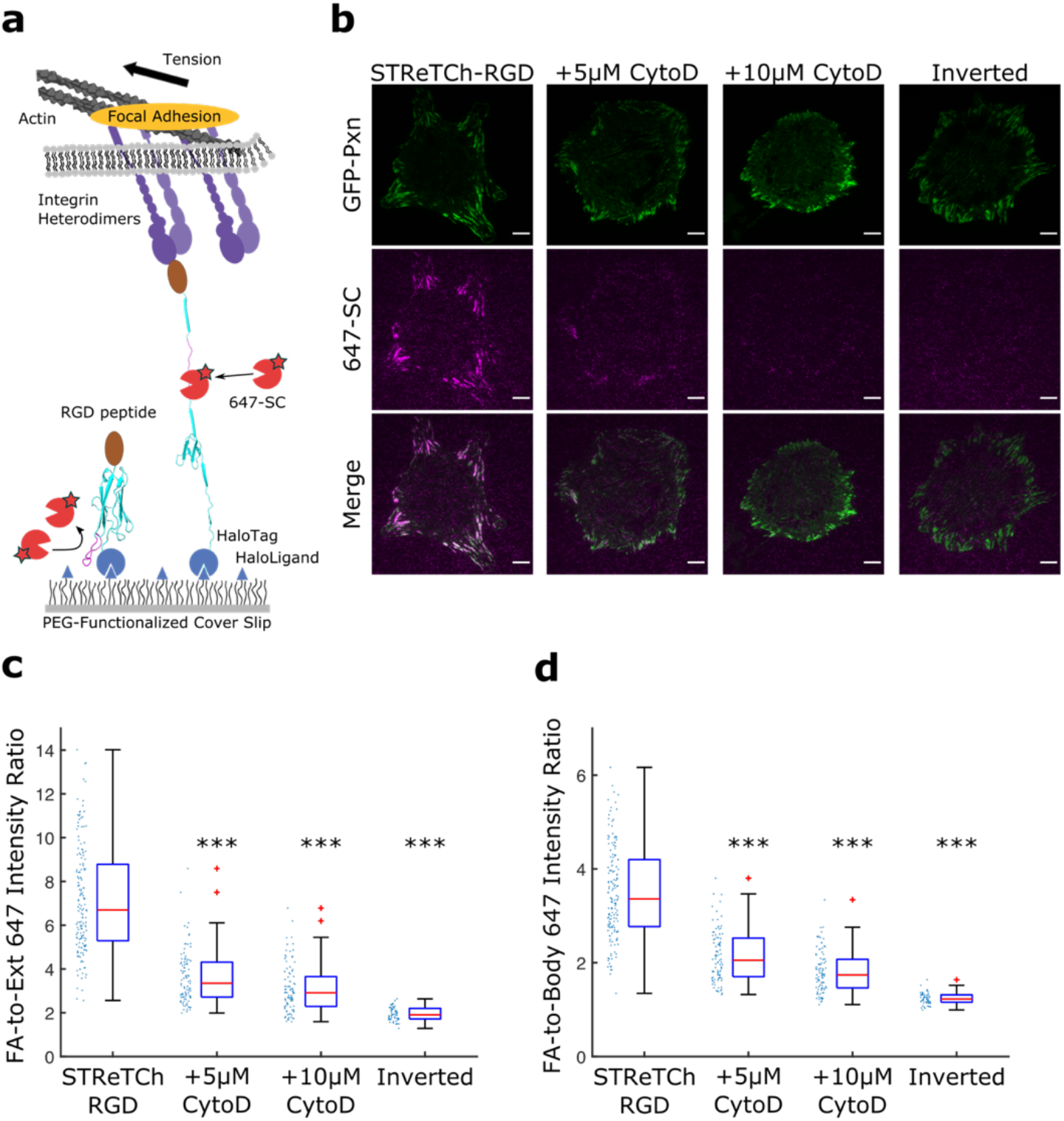
STReTCh detects cell adhesive forces across integrin-RGD bonds. **(a)** Experimental setup for extracellular force assay with STReTCh. Cells are seeded on a surface functionalized with STReTCh fused to an RGD ligand. Sensors under tension are visualized using SpyCatcher labeled with Alexa Fluor 647. **(b)** Representative images of GFP-paxillin and 647-SC for HFFs adhering to STReTCh-RGD in the absence and presence of Cytochalasin D, and for HFFs adhering to an inverted, force-inert construct. Scale bars = 10 μm. **(c)** Ratio of the average 647-SC intensity within FAs compared to background signal outside of cells for conditions described in (b). *** denotes p < 0.001 compared to STReTCh-RGD (two-tailed Mann-Whitney). Red lines indicate medians. Top and bottom of blue boxes indicate 75^th^ and 25^th^ percentiles, respectively, black bars indicate range, excluding outliers, and outliers are plotted as red plus signs. All subsequent boxplots follow this convention. Intensities for individual cells are plotted to the left of each bar. **(d)** Ratio of average 647-SC intensity within focal adhesions and average intensity outside of focal adhesions but underneath cell bodies. *** denotes p < 0.001 compared to STReTCh-RGD (two-tailed Mann-Whitney). *N* = 162 cells for STReTCh-RGD, 95 for 5 μM CytoD, 95 for 10 μM CytoD, and 59 for the inverted sensor. Data for each condition are pooled from a minimum of 3 independent experiments.

We observed strong colocalization of 647-SC and GFP-Pxn, indicative of 647-SC recruitment to FAs. As a control for staining in the absence of force, we seeded HFFs on a substrate coated with a force-inert variant of the fusion protein in which the STReTCh domain is C-terminal to the RGD peptide. Cells seeded on this variant can adhere to the surface through the RGD ligand, but tension is not transduced through the STReTCh domain^8^. As anticipated, we observed minimal staining in the force-inert control (**Fig. 3b-d**). Quantification of the average Alexa Fluor 647 fluorescence intensity inside of focal adhesions compared to the intensity outside of the cell yielded a signal-to-background ratio of ∼7:1 (**Fig. 3c, Supp. Fig. 3**). Overall signal intensities increased with increasing concentrations of 647-SC added (**Supp. Fig. 4**). Staining of STReTCh-RGD for HFFs imaged live, without fixation, yielded similar results when imaged using either TIRF (**Supp. Fig. 5**) or epifluorescence microscopy (**Supp. Fig. 6**). The 647-SC signal at FAs was reduced in cells treated with Cytochalasin D, which disrupts actin-mediated cytoskeletal tension in a concentration-dependent manner (**Fig 3c, d, Supp. Fig. 3**). These observations indicate that STReTCh robustly detects extracellular forces across individual integrins.

**Figure 4.**
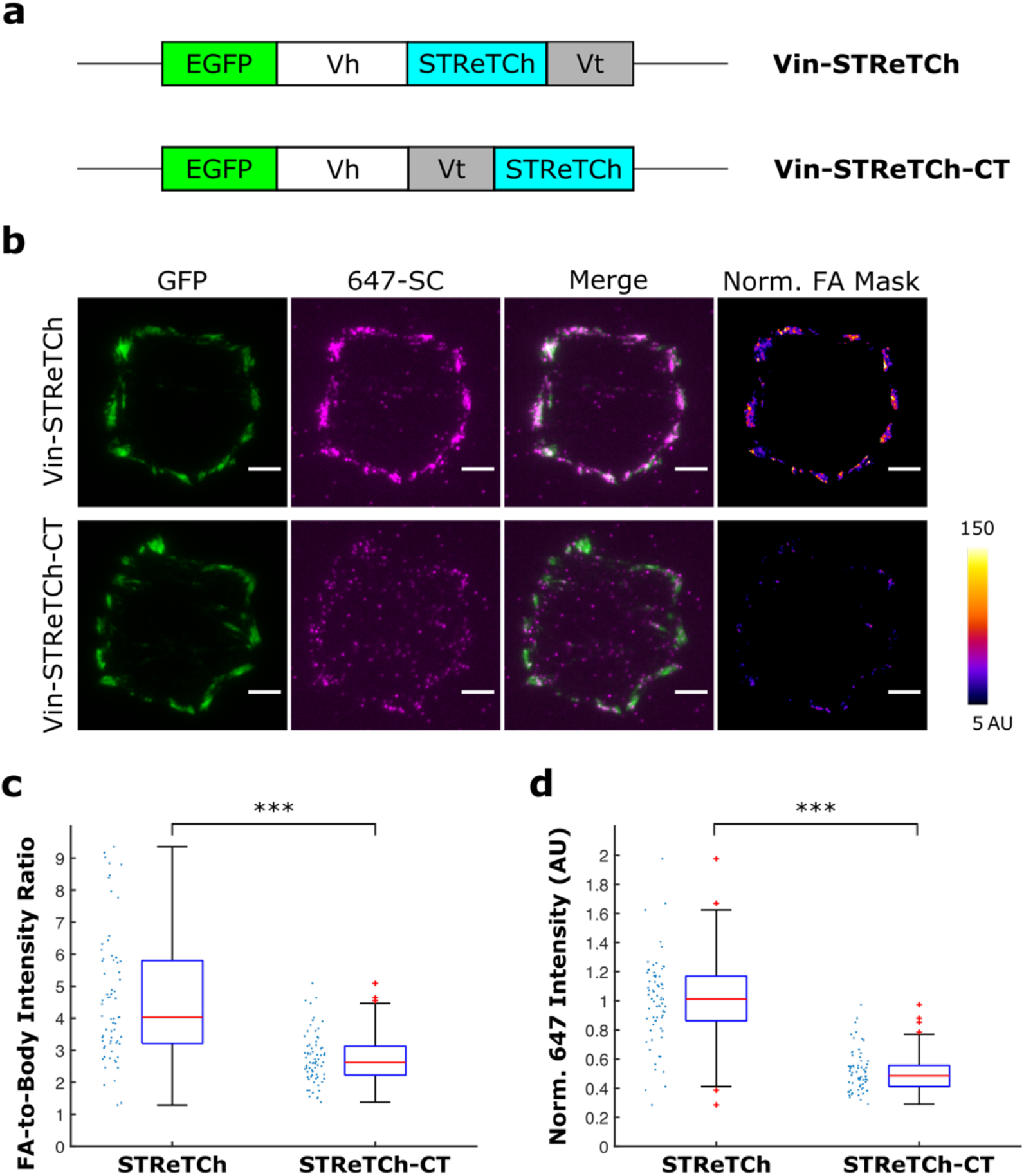
STReTCh detects intracellular tension across vinculin. **(a)** Constructs for vinculin-STReTCh (Vin-STReTCh) and a C-terminal force-inert control (Vin-STReTCh-CT). Vh denotes vinculin head. Vt denotes vinculin tail. **(b)** Representative images of GFP, 647 intensity, and 647 intensity normalized by GFP within FAs for MEFs expressing Vin-STReTCh and Vin-STReTCh-CT and fixed and stained with 647-SC. Scale bars, 5 μm. **(c)** Ratios of 647-SC intensity within FAs to the average intensity outside of FAs but within the cell boundary for MEFs expressing Vin-STReTCh and Vin-STReTCh-CT. **(d)** Quantification of 647 intensity normalized by GFP signal within FAs for Vin-STReTCh vs Vin-STReTCh-CT MEFs. *N* = 68 cells for STReTCh and 70 for STReTCh-CT, and data for each condition are pooled from 3 independent experiments. *** denotes p < 0.001 by two-tailed Mann-Whitney.

### Measuring intracellular forces across vinculin using STReTCh

After validating the functionality of STReTCh as an extracellular force sensor, we asked whether STReTCh could be genetically encoded to detect intracellular tension. In the experiments described above, 647-SC was added to cells after light fixation, but prior to permeabilization. Control experiments revealed some nonspecific sticking of 647-SC in fully fixed and permeabilized cells, including low levels of recruitment to FAs that presumably reflects the interaction of 647-SC with one or more cytoskeletal proteins (**Supp. Fig. 7**). We took advantage of the covalent nature of the SpyTag-SpyCatcher interaction and incorporated more stringent post-staining wash steps with EDTA and guanidine-HCl (Gdn-HCl), which reduced this nonspecific staining (**Supp. Fig. 8**).

To demonstrate the ability of STReTCh to measure intracellular tension, we inserted STReTCh into vinculin, a protein that reinforces connections between cell adhesion complexes and the actin cytoskeleton^3, 22^. Previous measurements found that vinculin on average bears 2-3 pN of force at focal adhesions in mouse embryonic fibroblasts (MEFs)^3^. In analogy to previous FRET-based tension sensors^3^, we inserted the STReTCh sensor between the vinculin head and tail domains in a flexible linker region known to be mechanosensitive. We further tagged this fusion protein with EGFP to quantify the expression levels and localization of the sensor construct, and denote this construct Vin-STReTCh (**Fig. 4a**). As a force-inert control, we placed STReTCh at the C-terminus of the GFP-vinculin fusion protein (Vin-STReTCh-CT) such that STReTCh would not bear tension (**Fig. 4a**). We stably transduced vinculin-null MEFs with these vinculin constructs and seeded these MEFs on fibronectin-coated coverslips. The MEFs were then fixed, permeabilized, blocked with bovine serum albumin, and incubated with 647-SC. Note that previous measurements show that measurable cytoskeletal tension persists up to 1 hour or more after fixation and permeablization^5, 17^, a timescale that is compatible with labeling with 647-SC. Finally, cells were washed with 2 mM EDTA and 3 M Gdn-HCl to remove residual, nonspecifically adsorbed 647-SC.

TIRF images of the fixed and stained Vin-STReTCh and Vin-STReTCh-CT MEFs are presented in **Figure 4**. The 647-SC signal was increased in the FAs of Vin-STReTCh MEFs as compared to Vin-STReTCh-CT MEFs, indicative of force-dependent exposure and labeling of the SpyTag sequence (**Fig. 4b-d, Supp. Fig. 8, Supp. Fig. 9**). Both the absolute and GFP-normalized 647-SC signal within FAs indicated higher degrees of 647-SC staining in Vin-STReTCh compared to Vin-STReTCh-CT (**Fig. 4b-d**). Expression levels of vinculin-STReTCh fusion proteins were relatively well-matched in both populations of cells, and 647-SC signal was higher in Vin-STReTCh MEFs irrespective of expression levels of the vinculin fusion proteins at FAs (**Supp. Fig. 10, 11**). Force across vinculin in Vin-STReTCh MEFs could also be visualized by 647-SC when images were acquired using epifluorescence illumination, though signal-to-background was lower and stringent wash steps with Gdn-HCl and EDTA seemed to be more critical for reducing non-specific sticking (**Supp. Fig. 12**). Overall, the increased focal adhesion staining in Vin-STReTCh MEFs compared to Vin-STReTCh-CT MEFs demonstrates that STReTCh can be inserted into proteins of interest to provide a readout of intracellular tension.

## DISCUSSION

In summary, we engineered a 11 kDa tension sensing protein module, termed STReTCh, by inserting SpyTag into the human titin I10 domain such that SpyTag is labeled when the I10 domain undergoes force-dependent unfolding. We used magnetic tweezers to characterize the unfolding behavior of STReTCh, which is sensitive to forces in the single pN range. STReTCh undergoes reversible unfolding at approximately 1 pN of applied force, making it among the most sensitive molecular tension sensors of which we are currently aware. We demonstrated the ability of STReTCh to detect biologically relevant forces associated with cell-ECM adhesion, both intra- and extracellularly. Used in an extracellular setting, STReTCh robustly reports on forces associated with integrin-ECM adhesion. Although SpyCatcher appeared to suffer from a low level of non-specific binding inside the cell, the covalent nature of the SpyCatcher-SpyTag bond enabled wash steps in the protocol that largely ablated non-covalent, non-specific interactions.

Despite its useful properties, the current STReTCh sensor does have limitations. The background signal due to nonspecific interaction of SpyCatcher with intracellular proteins currently limits signal-to-noise. Optimization of SpyCatcher to reduce background interactions or implementation of alternative tag/binder pairs may address this limitation. Unlike some FRET-based sensors^3, 5, 6, 23^, STReTCh is not meant to yield quantitative force measurements. However, quantitative force measurements are arguably required in only a minority of situations. For this reason, we anticipate that the ability of STReTCh to report on forces above a biologically meaningful force of ∼2 pN will make it a useful addition to the available suite of tools for force measurement. Finally, the current version of STReTCh is not compatible with intracellular, live-cell measurements, though it is sufficiently sensitive to measure residual intracellular forces in fixed cells, which persist approximately 1 hour after fixation^5, 17^. This limitation may potentially be addressed by versions in which the interaction of SpyCatcher and SpyTag (or other tag/binder pair) is reversible rather than covalent. Nevertheless, the experiments in this study demonstrate the utility of STReTCh as a general strategy for streamlined detection of molecular-scale forces in biological systems.

## ACKNOWLEDGEMENTS

B.L.Z is supported by the Stanford ChEM-H Chemistry/Biology Interface Predoctoral Training Program (NIH 5T32GM120007) and the NSF Graduate Research Fellowship Program (DGE-1656518). V.T.V. is supported by the Stanford Medical Scientist Training Program (NIH T32GM007365) and the NIH NIDDK (1F30DK124985). A.R.D. acknowledges the HHMI (Faculty Scholar Award) and the NIH (R35GM130332). We are grateful to current and former members of the Dunn lab—particularly Dr. Magnus Bauer, Dr. Steven Tan, and Dr. Cayla Miller—for discussion and comments on the manuscript.

## CONTRIBUTIONS

B.L.Z. and A.R.D. conceived the study and designed experiments. B.L.Z. and V.T.V. designed and built magnetic tweezers and implemented methods for magnetic tweezers data analysis. B.L.Z performed all experiments. B.L.Z. and A.R.D. wrote and prepared the manuscript, with input from V.T.V.

