## SupplementaryInfo for "STReTCh: a strategy for facile detection of mechanical forces across proteins in cells"

### MATERIALS AND METHODS

#### *DNA Constructs*

The STReTCh tension sensing module was designed by taking the titin I10 domain (residues 2880 to 2967 of *H. sapiens* titin, Q8WZ42 on UniProt) and inserting the 13-amino acid SpyTag (AHIVMVDAYKPTK) between residues 2956 and 2957. The amino acid sequence for the STReTCH module linked to a C-terminal 6xHis tag through a triglycine linker is:

METLHITKTMKNIEVPETKTASFECEVSHFNVPSMWLKNGVEIEMSEKFKIVVQGKLHQLIIMNTSTEDS
AEYTFVCGAHIVMVDAYKPTKNDQVSATLTVTGGGHHHHHH

For melting temperature characterization, the DNA encoding this construct was cloned into the pJ414 expression vector (DNA 2.0) by Epoch Life Sciences Inc. (Missouri City, TX). DNA encoding the wild type *H. sapiens* titin I10 domain was cloned by removing the SpyTag using Q5 site-directed mutagenesis (New England Biolabs) following manufacturers' instructions with the primers 5'-AACGATCAGGTGAGCGCG-3' and 5'- GCCGCACACAAAGGTATATTC-3'.

For magnetic tweezers force spectroscopy measurements, DNA encoding a protein composed of STReTCh fused to HaloTag at the N-terminus and a YbbR tag[1], TEV cleavage sequence, EGFP, and 6xHis tag at the C-terminus was cloned into pJ414 using custom cloning from Epoch Life Sciences Inc. Elastin-like polypeptide for functionalization of coverslips was expressed from pET28a-GGG-ELP(120nm)-Cys, a kind gift from Dr. Hermann Gaub (Addgene plasmid #91572 ; http://n2t.net/addgene:91572 ; RRID: Addgene_91572). A plasmid encoding the phosphopantetheinyl transferase Sfp from *B. subtilis* (Sfp synthase) was a kind gift from Dr. Michael Burkart (Addgene plasmid # 75015 ; http://n2t.net/addgene:75015 ; RRID: Addgene_7015).

For measurement of extracellular forces, the DNA encoding STReTCh-RGD was constructed from a previously described RGD force sensor construct encoding an N-terminal HaloTag, followed by a flagelliform-based tension sensing domain and a C-terminal RGD ligand[2-4]. STReTCh was inserted in place of the flagelliform tension sensing domain using Gibson Assembly (New England Biolabs). The amino acid sequence of STReTCh-RGD is:

MGSEIGTGFPFDPHYVEVLGERMHYVDVGPRDGTPVLFLHGNPTSSYVWRNIIPHVAPTHRSIAPDLIG
MGKSDKPDLGYFFDDHVRFMDAFIEALGLEEVVLVIHDWGSALGFHWAKRNPERVKGIAFMEFIRPIPT
WDEWPEFARETFQAFRTTDVGRKLIIDQNVFIEGTLPMGVVRPLTEVEMDHYREPFLNPVDREPLWRF
PNELPIAGEPANIVALVEEYMDWLHQSPVPKLLFWGTPGVLIPPAEAARLAKSLPNAKAVDIGPGLNLLQ
EDNPDLIGSEIARWLSTLEISGGAGEF**METLHITKTMKNIEVPETKTASFECEVSHFNVPSMWLKNGVE
IEMSEKFKIVVQGKLHQLIIMNTSTEDSAEYTFVCGAHIVMVDAYKPTKNDQVSATLTVT**SENLYFEQG
TVYAVTGRGDSPASSAAHHHHHH

The force-inert, inverted RGD sensor was cloned into pJ414 by Epoch Life Sciences Inc. and has the following amino acid sequence:

MGSEIGTGFPFDPHYVEVLGERMHYVDVGPRDGTPVLFLHGNPTSSYVWRNIIPHVAPTHRSIAPDLIG
MGKSDKPDLGYFFDDHVRFMDAFIEALGLEEVVLVIHDWGSALGFHWAKRNPERVKGIAFMEFIRPIPT
WDEWPEFARETFQAFRTTDVGRKLIIDQNVFIEGTLPMGVVRPLTEVEMDHYREPFLNPVDREPLWRF
PNELPIAGEPANIVALVEEYMDWLHQSPVPKLLFWGTPGVLIPPAEAARLAKSLPNAKAVDIGPGLNLLQ
EDNPDLIGSEIARWLSTLEISGGAGEFTVYAVTGRGDSPASSAAGGG**METLHITKTMKNIEVPETKTAS
FECEVSHFNVPSMWLKNGVEIEMSEKFKIVVQGKLHQLIIMNTSTEDSAEYTFVCGAHIVMVDAYKPT
KNDQVSATLTVT**SENLYFEQGHHHHHH

DNA encoding a truncated, minimal SpyCatcher (SCMin) with a cysteine for thiol-maleimide labeling was cloned from pDEST14-SpyCatcher (a gift from Mark Howarth, Addgene plasmid # 35044 ;

http://n2t.net/addgene:35044 ; RRID:Addgene_35044). A cysteine was introduced through a Y8C mutation downstream of the N-terminal 6xHis Tag using QuikChange II XL site-directed mutagenesis (Agilent). The full-length SpyCatcher was then truncated using Gibson Assembly to remove amino acids 21-43 and the C-terminal 9 amino acids to give the final amino acid sequence for the minimal SpyCatcher (SCMin):

HHHHHHDCDIPTTENLYFQGDSATHIKFSKRDEDGKELAGATMELRDSSGKTISTWISDGQVKDFYLYP
GKYTFVETAAPDGYEVATAITFTVNEQGQVTVNG

DNA constructs for Vin-STReTCh and Vin-STReTCh-CT were cloned from a PiggyBac plasmid containing *G. gallus* vinculin downstream of the hEF1$\alpha$ promoter (pJ509-02, DNA 2.0). EGFP linked to a GS linker (SGLGGSGGGGSGGGGSGG) was fused at the N-terminus of vinculin. STReTCh was inserted either after amino acid 883 of vinculin (between the head and tail domains) or at the C-terminus as illustrated in Fig. 3. These fusion proteins were cloned into the PiggyBac vector downstream of the hEF1$\alpha$ promoter by Epoch Life Sciences Inc. to give EGFP-Vinculin-STReTCh(-CT). All constructs were verified by sequencing.

***Protein expression, purification, and labeling***

All proteins were expressed in BL21(DE3) chemically competent *Escherichia coli* and purified using Ni-NTA chromatography, except for the ELP. Cultures were induced at an optical density of 0.6 with 1mM isopropyl-β-D-thiogalactopyranoside and grown overnight at 18 ºC. The cultures were then spun down at 6000×*g* for 15 minutes, and bacterial pellets were resuspended in lysis buffer (50 mM sodium phosphate, 300 mM NaCl, and 10 mM imidazole, pH 8) with a cOmplete EDTA-free protease inhibitor cocktail (11873580001, Roche), 70 μg/mL lysozyme (90082, ThermoFisher), and 7 U/mL DNase I (04536282001, Roche). The resuspended cells were rotated end-over-end for 45 min at 4 °C, lysed with a tip sonicator, and spun at 12,000×*g* for 30 min. The supernatant was incubated with 1.5 ml of nickel–nitrilotriacetic acid HisPur Resin (ThermoFisher) for every 500 mL of culture and rotated end-over-end at 4 °C for 2 hours. The solution was then packed into a gravity column, washed three times with 5 ml of wash buffer (50 mM sodium phosphate, 300 mM NaCl, and 20 mM imidazole, pH 7.4 with 2 mM β-mercaptoethanol; BME), and protein was eluted by incubating the resin bed with 1 mL of elution buffer (50 mM sodium phosphate, 300 mM sodium chloride, and 250 mM imidazole, pH 7.4 and 2 mM BME) for 5 minutes, four times. The eluate was collected, concentrated, and buffer-exchanged into storage buffer (1× phosphate-buffered saline, 2 mM BME) using Amicon centrifugal filter units (MilliporeSigma) of the appropriate molecular-weight cutoff, then flash-frozen and stored at −80 °C. For magnetic tweezers experiments, the HaloTag-STReTCh-YbbR-GFP fusion protein was further purified using anion exchange chromatography with a 1 mL HiTrap Q column (Cytiva Life Sciences) in 50mM Tris, pH 8 supplemented with 2 mM BME using an AKTA Pure FPLC (GE Healthcare). FPLC purified protein was then buffer-exchanged into PBS + 2 mM BME before freezing. Eluates were characterized by SDS–polyacrylamide gel electrophoresis (SDS-PAGE), and the concentration was determined by ultraviolet–visible (UV-Vis) spectroscopy.

For magnetic tweezer experiments, ELP was expressed in BL21(DE3) *E. coli* and purified using a thermoprecipitation and redissolution strategy described previously[5], then buffer exchanged into 50 mM HEPES prior to flash freezing. Biotinylated STReTCh was generated by labeling HaloTag-STReTCh-YbbR-GFP with biotin-PEG3-CoenzymeA (SiChem GmbH, Bremen, Germany). 5 μM HaloTag-STReTCh-YbbR-GFP was incubated with 0.5 μM Sfp synthase, 10 μM biotin-PEG3-CoenzymeA, and 10 mM $MgCl_2$ in PBS overnight rotating at 4 ºC to give HaloTag-STReTCh-Biotin-GFP. The reaction was then buffer exchanged into PBS + 2 mM BME using a 7 kDa Zeba desalting column to remove excess biotin-PEG3-CoenzymeA, flash-frozen, and stored at -80 ºC for later use.

For in-cell experiments, Alexa Fluor 647 labeled SpyCatcher (647-SC) was generated by first further purifying SCMin using size exclusion chromatography on an AKTA Pure FPLC with a Sephadex 75

Increase 10/300 GL (Cytiva Life Sciences) column in PBS + 2 mM BME. Purified SCMin was buffer-exchanged into buffer containing 100 mM phosphate, 150 mM NaCl and 1 mM EDTA using three 7 kDa Zeba desalting columns (ThermoFisher) in series to remove excess BME. 100 μL of 116 μM desalted SpyCatcher was then incubated with excess tris(2-carboxyethyl)phosphine (TCEP) for 1 hour at room temperature, followed by 2 equivalents of Alexa Fluor 647 $C_2$-maleimide (ThermoFisher, A20347), rotating at 4 ºC overnight. The reaction was then quenched by adding 20 μL of PBS + 2 mM BME. Excess free dye was removed using 2 Sephadex G-25 PD MiniTrap desalting columns (Cytiva Life Sciences) in series, using gravity-driven flow. Labeling efficiency was 72% as measured by UV-Vis. The labeled 647-SC was flash-frozen at -80 ºC and stored in PBS until later use.

#### *Melting temperature characterization*

STReTCh or titin I10 were mixed with SYPRO Orange dye (ThermoFisher) at a final concentration of 5 μM protein and 5x SYPRO Orange. 3 replicates of 35 μL of the protein-dye mixture were loaded into a 96-well PCR plate (ThermoFisher) which was spun down in a swinging bucket centrifuge prior to measurement. SYPRO Orange fluorescent signal was measured at temperatures from 25 ºC to 95 ºC at 0.5 ºC increments using a StepOnePlus Real-Time PCR instrument (ThermoFisher).

#### *Magnetic tweezer construction and calibration*

A magnetic tweezer was constructed on a Nikon Ti-E microscope equipped with a motorized 3-axis piezo stage (Mad City Labs). Two 6 mm-diameter cylindrical N52 magnets (K&J Magnets), arranged as in Supp. Fig. 2, were affixed to a custom-fabricated aluminum bracket, which was attached to a 0.5”-travel XYZ translation stage equipped with a motorized Z-axis actuator (Thorlabs, Newton, New Jersey, USA). The stage position was homed before measurement by stepping the Z-actuator until the magnet bracket just touched the top of the sample. Measurements were performed while the Nikon Perfect Focus feedback system was turned on.

Samples were illuminated from the objective side via a mint green LED (Thorlabs, MINTL5) reflected into the back focal plane of a 100x Plan-Apo TIRF objective using a polarizing beamsplitter (Edmund Optics). Reflected, cross-polarized light was transmitted through the beamsplitter and collected by a sCMOS camera (Hamamatsu Orca Flash v4). Bead images were collected using a custom-written LabView script or MicroManager 1.4 software[6].

Magnetic tweezers force calibration was performed as described previously[6] using transverse fluctuations of a magnetic bead tethered to a glass coverslip via lambda-phage DNA (New England Biolabs).

#### *Magnetic tweezers force spectroscopy*

Coverslips functionalized with HaloLigand-ELP were prepared similarly to a previously described protocol[2-4]. ELPs were first functionalized at the N-terminus with HaloLigand by reacting 300 μM ELP with 3 mM HaloTag Succinimidyl Ester (O4) Ligand (Promega, P6751) in 50 mM HEPES, pH 7.5 for 4 hours at room temperature. During this reaction, glass coverslips (Fisher Scientific, No. 12-544-14) were cleaned and silanized as described previously[2-4] to generate a nucleophilic, amine-functionalized substrate. Sulfosuccinimidyl 4-(N-maleimidomethyl)cyclohexane-1-carboxylate (ThermoFisher, A39268) (Sulfo-SMCC) was diluted to 10 mM in 50 mM HEPES, pH 7.5, and a coverslip “sandwich” was formed by adding 90 μL of the Sulfo-SMCC solution between a pair of silanized coverslips. These coverslip sandwiches were incubated at room temperature in a humid environment for 1 hour to generate maleimide-functionalized coverslips. Coverslips were washed by dipping in a beaker of MilliQ water to remove excess Sulfo-SMCC, then gently dried under a stream of nitrogen.

After the 4 hour incubation was complete, the HaloLigand-ELP reaction was quenched by adding Tris buffer (pH 7) to a final concentration of 5 mM for 15 minutes. In preparation for functionalizing the ELP to the coverslip using cysteine-maleimide chemistry, excess TCEP was added to the reaction mixture and incubated for 30 minutes at room temperature. The reaction was then buffer exchanged into ELP conjugation buffer (50 mM phosphate, 50 mM NaCl, 10 mM EDTA, pH 7.2) using two 7 kDa Zeba desalting columns in series. Coverslip sandwiches using the maleimide-functionalized coverslips were incubated with 90 µL of the ELP solution for 2 hours at room temperature in a humid environment. Coverslips were then washed by dipping in MilliQ water as described above, and were sufficiently dry upon slow removal of the coverslips from the water due to the relatively hydrophobic surface properties of the ELP-functionalized surface. Finally, ELP-functionalized coverslip sandwiches were incubated with 100 µL of 10 mM L-cysteine for 1 hour at room temperature in a humid environment to quench remaining free maleimides. These coverslips were then washed by dipping in water and were sufficiently dry for storage without additional drying steps. Coverslips were stored in vacuum-sealed bags at room temperature for up to several weeks, or at -20 ºC for longer term storage.

Non-magnetic 2.0 µm polystyrene beads (Polybeads, Polysciences) were non-specifically attached to the HaloLigand-ELP coverslips to use as reference beads to account for drift during magnetic tweezer measurements. 1 drop (~50 µL) of polystyrene bead stock suspension was diluted in 1.5 mL ethanol, and 100 µL was spin-coated onto the coverslip at 3500 rpm for 40 s. The coverslip was then baked at 75 ºC for 10 minutes, then washed by dipping in MilliQ water to remove non-adhered polystyrene beads.

3-well coverwells (Grace Biolabs) of ~100 µL well volumes were attached to HaloLigand-ELP functionalized coverslips. Wells were surface passivated by treatment for 1 hour at room temperature with 110 µL of a 1% (w/v) solution of casein, diluted from a 5% stock (Sigma-Aldrich, C4765). Wells were then washed twice with 200 µL PBS. 110 µL of HaloTag-STReTCh-Biotin-GFP was added to the wells at concentrations ranging from 100 pM to 1 nM and incubated at room temperature for 30 minutes to allow for covalent attachment to the coverslip. Wells were washed twice with 200 µL PBS to remove unbound protein.

Streptavidin-coated superparamagnetic beads (Dynabeads M-270 Streptavidin, ThermoFisher, 65305) were washed by diluting the stock 30x in PBS + 0.1% Tween-20 and vortexing the diluted bead suspension for 20 s. Beads were pulled down using a magnet, and washed twice more with PBS + 0.1% Tween-20. Beads were then resuspended in PBS + 0.1% Tween-20 + 1% casein at a 30x dilution from the stock concentration and equilibrated with the buffer for 15 minutes. 110 µL of this bead suspension was added to the HaloTag-STReTCh-Biotin-GFP functionalized wells and incubated at room temperature for 25 minutes. After bead incubation, magnetic tweezers were brought above the wells to apply sub-pN forces to the samples to remove a majority of the beads not tethered to the biotinylated STReTCh construct, and the wells were washed twice with 200 µL PBS. This detachment and wash process was repeated twice before making magnetic tweezers measurements. Minutes-long movies of streptavidin beads tethered to HaloTag-STReTCh-Biotin-GFP surface were then acquired at frequencies ranging from 2.5 Hz to 50 Hz, with a 20 ms exposure time, at forces ranging from sub-pN to ~8 pN.

***Magnetic tweezers data analysis***

As described previously[7], in separate calibration experiments, a nonspecifically adhered bead was imaged while the piezo stage was stepped in increments of 5 nm to generate a reference image lookup table (LUT). These LUTs were generated for both magnetic and non-magnetic beads and were used to estimate the *z*-position of query bead images. The analysis pipeline is schematized in Supp. Fig. S2. Briefly, the two-dimensional Fourier transform of the query bead image was calculated, and the radial profile extracted. The bead position was then estimated in one of two ways. In one method, the Pearson correlation coefficient *r* between the radial profile of the query image and each image in the LUT were calculated, and the LUT position that maximized *r* was computed by quadratic interpolation.

In the second method, the one-dimensional Fourier transform of the LUT radial profiles was computed. The phase at a high-amplitude frequency was then plotted against the bead height corresponding to the LUT slice, and a third- to fifth-order polynomial was fit to this curve. This polynomial fit was then used to directly estimate height from the Fourier transform of the query image. Images were analyzed with both methods and found to produce similar results, reducing the chance that measured bead heights were analysis artifacts.

To correct for drift during measurement, the *z*-position trace of either a non-magnetic polystyrene bead or occasionally an immobile, non-specifically adhered magnetic bead was subtracted from the trace of the bead of interest. To characterize the unfolding behavior of the STReTCh domain, we identified traces in which the bead of interest exhibited steplike changes in length of ~10-15 nm. We further filtered these traces based on the height extension behavior to validate the presence of ELP extension, which has a contour length of ~120 nm[5, 7], and by confirming that the bead exhibited *xy*-fluctuations consistent with the expected Brownian motion under pN forces based on the Equipartition Theorem[8]. Steps in traces that met the quality control criteria were manually identified, and datapoints for each trace were designated as either folded or unfolded based on the localized, reference-subtracted *z*-position to calculate the fraction of the trace that the molecule spent in each of these two states. The folded fraction was then plotted in Figure 1b.

Magnetic tweezers data for the folding and unfolding of STReTCh was fit to the Bell-Evans kinetic model[9], which gives a sigmoidal fit for the folded fraction as illustrated in Figure 1b. The equilibrium between the folded species $P_f$ and unfolded species $P_u$ is:

$$P_f \underset{k_f}{\overset{k_u}{\rightleftharpoons}} P_u$$

where $k_u$ and $k_f$ are the first-order rate constants for folding and unfolding, respectively, and are dependent on the applied force $F$:

$$k_u = k_u^0 e^{\frac{\Delta x_u F}{k_B T}}$$
$$k_f = k_f^0 e^{\frac{\Delta x_f F}{k_B T}}$$

Here, $k_u^0$ and $k_f^0$ are force-independent rates, $\Delta x_u$ and $\Delta x_f$ represent distances to the transition state from the folded and unfolded states, $k_B$ is Boltzmann's constant, and $T$ is the temperature. At equilibrium, the folding and unfolding rates are equal, so concentrations of the folded and unfolded species satisfy

$$k_u P_f = k_f P_u$$

and the folded fraction is given by

$$\frac{P_f}{P_f + P_u} = \frac{\frac{k_f P_u}{k_u}}{\frac{k_f P_u}{k_u} + P_u} = \frac{1}{1 + \frac{k_u}{k_f}}$$

$$\frac{P_f}{P_f + P_u} = \frac{1}{1 + \frac{k_u^0}{k_f^0} e^{\frac{(\Delta x_u - \Delta x_f)F}{k_B T}}}$$

which varies sigmoidally with the applied force $F$.

We assume that the system is ergodic, such that the folded fraction of an ensemble molecules can be represented by the average fraction of time that individual molecules spend in the folded state (i.e. the data presented in Figure 1b). Consequently, we used the magnetic tweezers data to perform maximum likelihood estimation (MLE) fitting for 4 parameters: $k_u^0$, $k_f^0$, $\Delta x_u/k_BT$ and $\Delta x_f/k_BT$. For each trace, we define a probability of observing steps of the measured lengths by assuming that the waiting times for folding and unfolding are Poisson-distributed and that all folding and unfolding events are independent. Then, the probability density for observing a folded time step of duration $t_f$ or unfolded time step of duration $t_u$ is given by the expression

$$p(t_f) = k_u e^{-k_u t_f}$$
$$p(t_u) = k_f e^{-k_f t_u}$$

Additionally, because determining the waiting time of the last step in each trace is limited by when we stop observing the specific bead, we assign probabilities for these steps equal to the likelihood of a waiting time of *at least* as long as the duration that was observed. In other words, the probabilities associated with observing a folded time step $t_f$ or unfolded time step $t_u$ for the last steps in the trace are given by

$$p(t > t_f) = \int_{t_f}^{\infty} k_u e^{-k_u t} dt = e^{-k_u t_f}$$
$$p(t > t_u) = \int_{t_u}^{\infty} k_f e^{-k_f t} dt = e^{-k_f t_u}$$

Thus, for an observed trace with *n* observed steps (i.e. *n-1* transitions between folded and unfolded states) where each step *i* has a duration $t_i$, the probability of observing such a trace can be computed as

$$p(\{t_1, t_2 \dots t_n\}) = \left( \prod_{i=1}^{n-1} k_i e^{-k_i t_i} \right) e^{-k_n t_n}$$

Then the probability of observing a dataset of m traces given a set of Bell-Evans parameters is

$$p(\{trace_1, trace_2 \dots trace_m\}) = \prod_{i=1}^{m} \left[ \left( \prod_{j=1}^{n_i - 1} k_{j_i} e^{-k_{j_i} t_{j_i}} \right) e^{-k_{n_i} t_{n_i}} \right]$$

where each rate constant takes on the expression for $k_u$ or $k_f$ according to the type of step observed. The MLE fit for the 4 parameters are the values that maximize this probability expression. To calculate these estimates, we used the built-in MATLAB function fminsearch to minimize an objective function corresponding to the negative value of the natural logarithm of the probability expression above. These MLE fit parameters are tabulated in Table S1. We also used the logarithm of the probability expression as the log-likelihood function to calculate confidence intervals for MLE parameters using a profile likelihood approach[10], where a given parameter, generically *q*, is systematically varied about its optimal value, while all other parameters are optimized for each value of *q*. The resulting log-likelihood ratios asymptotically approach a $\chi^2$ distribution with one degree of freedom. The upper and lower bounds are thus determined by

$$2\{l(\hat{\theta}) - l(\hat{\theta}_{lower})\} = 2\{l(\hat{\theta}) - l(\hat{\theta}_{upper})\} = c_{1;1-\alpha}$$

Where $l$ is the log-likelihood function, $\hat{\theta}$ denotes the MLE for $q$, and $c_{1;1-\alpha}$ is the (1-$\alpha$)th quantile of the $\chi^2$ distribution with 1 degree of freedom. We solved this expression to find the upper and lower bounds for the 95% confidence interval, which are also tabulated in Table S1.

***Cell culture and transfection***

GFP-paxillin expressing HFFs were prepared from CCD-1070Sk HFFs (ATCC CRL-2091) as previously described[2-4]. Vin-STReTCh(-CT) MEFs were prepared by transfecting $vin^{-/-}$ MEFs, a gift from K. Rothenberg and B. Hoffman (Duke University). Both HFFs and MEFs were cultured in DMEM high-glucose medium (Gibco, catalog no. 21063-029) in the absence of phenol red and supplemented with 10% fetal bovine serum (FBS, Axenia Biologix LLC), sodium pyruvate (1 mM, Gibco), MEM nonessential amino acids (1×, Gibco), and penicillin/streptomycin (100 U/ml and 100 µg/ml, Gibco). Cells were cultured on tissue culture plastic and grown at 37 °C with 5% $CO_2$.

$Vin^{-/-}$ MEFs were stably transduced with EGFP-Vinculin-STReTCh or EGFP-Vinculin-STReTCh-CT using a Lonza P4 kit. Cells were trypsinized, pelleted, resuspended in medium without FBS and penicillin/streptomycin, then counted. $5{\cdot}10^5$ cells were repelleted, then resuspended in a solution of 82 µL P4 nucleofector solution and 18 µL P4 supplement. 2.5 µg of DNA was added to the cells and gently flicked before transferring to a Lonza nucleofection cuvette. Cuvettes were placed in a Lonza 4D-Nucleofector system and program C2167 (for MEFs) was used. 500 µL of warm medium was added to the cuvette to transfer the cells to a six-well plate with medium equilibrated at 37 °C using a pipette bulb without pipetting up and down. 24 hours after transfection, MEFs with the stably integrated constructs were selected for using puromycin at increasing concentrations (1-2 µg/mL) over the course of 5 days, then sorted into populations using fluorescence-assisted cell sorting based on GFP to enrich for construct-expressing cells and isolate populations of Vin-STReTCh and Vin-STReTCh-CT MEFs with similar levels of protein expression.

***TIRF Microscopy***

Total internal reflection fluorescence (TIRF) microscopy images were collected using an Apo TIRF 100× oil objective lens, numerical aperture 1.49 (Nikon) as described previously[2-4] and controlled using Micromanager[6]. Samples were excited with 473-nm OBIS laser (Coherent) or 635-nm (Blue Sky Research) lasers for GFP and Alexa Fluor 647, respectively. Emitted light passed through a quad-edge laser-flat dichroic with center/bandwidths of 405/60 nm, 488/100 nm, 532/100 nm, and 635/100 nm from Semrock Inc. (Di01-R405/488/532/635-25×36) and corresponding quad-pass filter with center/bandwidths of 446/37 nm, 510/20 nm, 581/70 nm, 703/88 nm band-pass filter (FF01-446/510/581/703-25). GFP and Alexa Fluor 647 images were taken through separate additional cubes stacked into the light path (GFP: 470/40 nm, 495 nm long-pass, and 525/50 nm; Alexa Fluor 647: 679/41 nm and 700/75 nm) and recorded on a Hamamatsu Orca Flash 4.0 camera.

***Extracellular force sensor measurements***

HaloLigand-PEG coverslips were prepared as described in the literature[2-4]. 3- and 4-well coverwells (Grace Biolabs) of ~100 µL well volumes were attached to the functionalized coverslip. 100 µL of 200 nM Halo-STReTCh-RGD or inverted control in PBS was flowed into each channel and incubated at room temperature for 45 minutes. Channels were washed with 200 µL PBS to wash out unbound sensor. GFP-Pxn HFFs were seeded onto the force sensor cover slips and incubated at 37 °C for 75-90 min to allow for cell spreading and adhesion onto the cover slip.

For live cell measurements, non-adhered cells were washed off with 200 µL media at 37 °C after spreading. Cells were then treated with 110 µL of 20 nM SC-647 in media and incubated at 37 °C for 20 minutes. After staining, cells were washed with 200 µL media at 37 °C and imaged using TIRF microscopy with the objective maintained at 37 °C. As illustrated in Supp. Fig. S5, we observed some movement of HFFs during live cell imaging. To better localize potential SC-647 signal in relation to focal adhesions, we treated cells with 110 µL 0.5% (v/v) paraformaldehyde, diluted in warm PBS from a 16% stock (Electron Microscopy Sciences) for 5 minutes at 37 ºC to lightly fix cells and reduce cell motion during imaging. After light fixation, cells were washed with 200 µL warm PBS, followed by 200 µL warm media and subsequently treated with 110 µL SC-647 at the desired concentration (10-40 nM) for 10 minutes at 37 ºC, washed with 200 µL warm media, then imaged using TIRF microscopy at 37 ºC to generate the data in Figure 2. For conditions in which Cytochalasin D (CytoD, Enzo Life Sciences) was added, 110 µL of CytoD diluted at the desired concentration in warm media was added 5 minutes prior to the light fixation procedure.

For fixed cell measurements (Supp. Fig. 7), HFFs were seeded on coverslips functionalized with Halo-STReTCh-RGD for 1 hr, after which nonadherent cells were washed off with 200 µL warm media followed by 200 µL PBS at 37 ºC. Cells were then fixed with 100 µL 4% PFA in PBS for 15 minutes and permeabilized with 100 µL 0.1% Triton-X in PBS for 10 minutes at room temperature. Cells were washed with 200 µL PBS after each step. Cells were then blocked with 100 µL 1% bovine serum albumin (BSA) in PBS for 20 minutes, then stained with 20 nM SC-647 in 1% BSA/PBS for 20 minutes. After staining, cells were washed 3 times with 200 µL PBS. Fixed and stained cells were then imaged with TIRF microscopy at room temperature.

***Intracellular force sensor measurements***

3- and 4-well coverwells (Grace Biolabs) were attached to cleaned coverslips. 100 µL of fibronectin (10 µg/mL, Corning) diluted in PBS was added to the well and incubated for 45 minutes at room temperature. Wells were washed with 200 µL PBS, then Vin-STReTCh(-CT) MEFs were seeded by adding 100 µL of a 300,000 cells/mL suspension in warm medium. Cells were allowed to adhere to the surface for 3 hours, after which, they were washed with warm media followed by warm PBS. Cells were then fixed with 100 µL 4% PFA in PBS for 15 minutes and permeabilized with 100 µL 0.1% Triton-X in PBS for 10 minutes at room temperature. Cells were washed with 200 µL PBS after each step. Cells were then blocked with 100 µL 1% BSA in PBS for 20 minutes, then stained with 10nM SC-647 in 1% BSA/PBS for 10 minutes. After staining, cells were either washed once with 200 µL PBS, then either 3x with 200 µL PBS, or washed once with 2 mM EDTA and twice with 3 M Gdn-HCl. Cells were incubated for 3 minutes with each wash solution, washed with 200 µL PBS between washes, and washed with 2x200 µL PBS after the final Gdn-HCl or PBS wash, for a total of 8 washes after staining. Fixed and stained cells were then imaged with TIRF microscopy at room temperature.

***Image analysis and quantification***

Image preparation and quantification of fluorescence intensities in specific regions of stained GFP-Pxn HFFs and Vin-STReTCh(-CT) MEFs was carried out in Fiji[11]. Images were background subtracted, using either rolling ball background subtraction for GFP signal or subtracting the average intensity of an empty, unstained sample for 647 signal. Focal adhesions were segmented by generating a mask from thresholding GFP signal using Fiji's built-in RenyiEntropy algorithm. The cell body was segmented by generating a mask from thresholding GFP signal using Fiji's built-in Triangle algorithm applied to a Gaussian blurred GFP image. The region of the image denoted as "Body" (Figs. 2 and 3, Supp. Figs. 3 and 9) is thus defined as the region outside the FA mask but within the cell body mask. The region of the image denoted as "Ext" is defined as the region outside the cell body mask.

To compute the 647 signal normalized to the expression of Vin-STReTCh(-CT) for intracellular force measurements, GFP images were background subtracted using rolling ball background subtraction, and the 647 signal within the cell body was background subtracted using the mean cell exterior value for each image. The background subtracted 647 image was divided by the background subtracted GFP image, and this resultant image was multiplied by a FA mask generated from the GFP signal using the method described above. The mean value within this mask was measured for each cell and reported as the normalized 647 intensity in Fig. 3d.

Metrics are normalized to the mean values within FAs for the force-bearing STReTCh condition when applicable to account for potential variations in imaging conditions (e.g. laser power, TIRF angle, etc.) between different experiments.

### SUPPLEMENTARY FIGURES AND TABLES

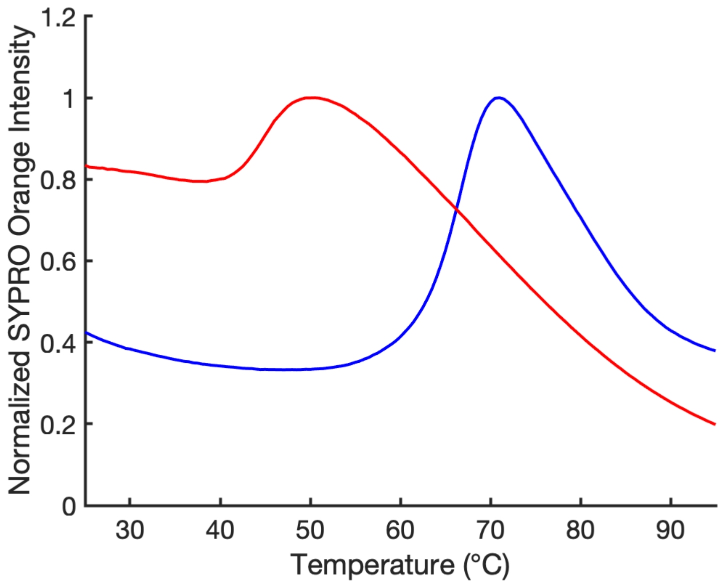


**Supplementary Figure 1. Melting curves of STReTCh and Titin I10**

Melting curves for the STReTCh module (*red*) and *H. sapiens* titin I10 domain (*blue*) as measured by SYPRO Orange dye fluorescence, normalized to the maximum intensity measured for each protein. Melting temperatures, measured as points of maximal slope, for STReTCh and titin I10 are 45 ºC and 66 ºC, respectively. The estimated melting temperature for titin I10 is consistent with previous measurements[1]. Curves are averages of triplicate measurements, with measurements performed on two separate days for STReTCh and once for titin I10.

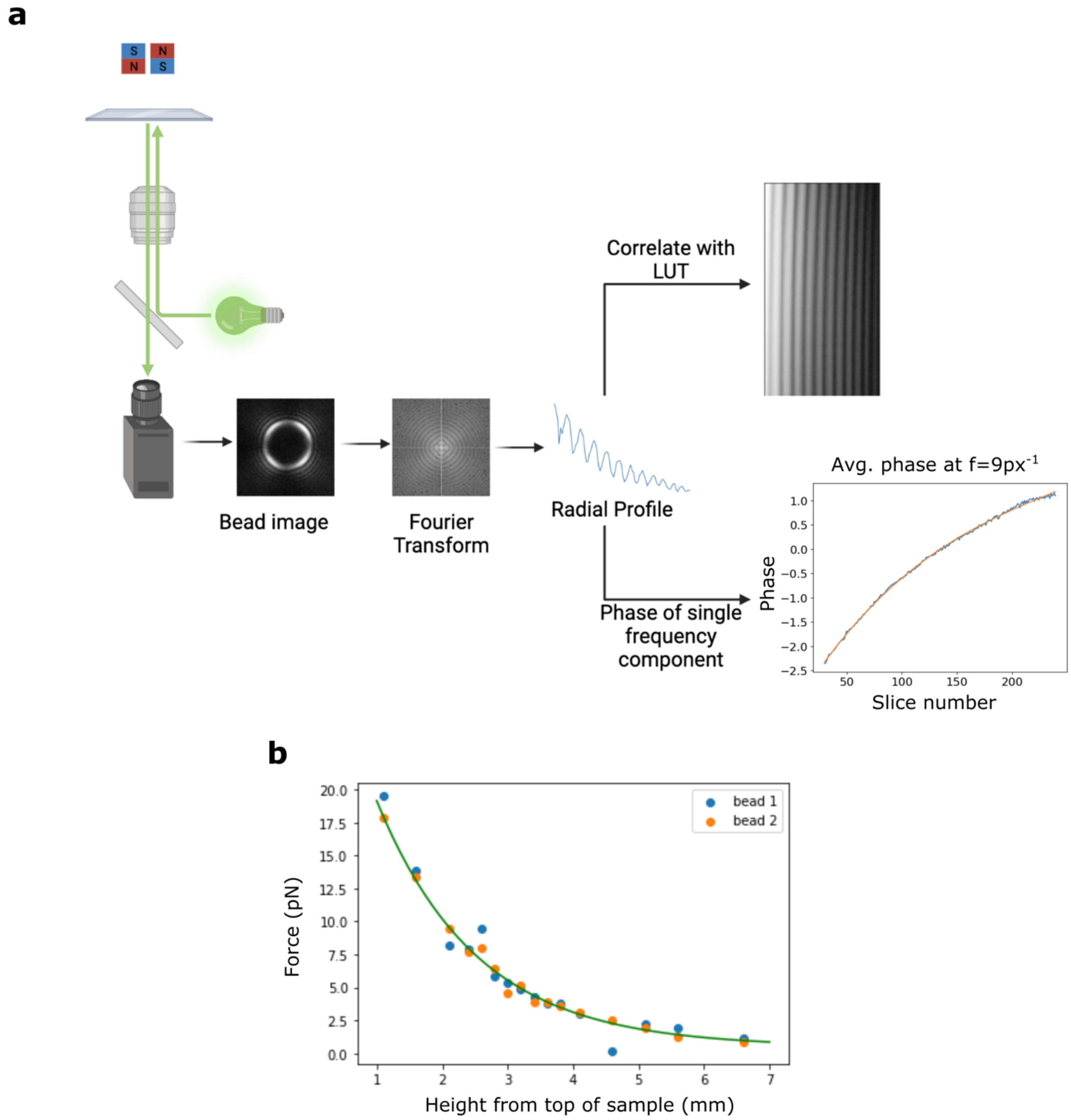


**Supplementary Figure 2. Magnetic tweezer setup, calibration, and data analysis and sample STReTCh unfolding traces**

**(a)** Schematic of magnetic tweezers setup and data analysis pipeline. **(b)** Force applied to M270 superparamagnetic beads as a function of distance of the magnetic tweezers from the sample. Relationship between force and distance follows a biexponential fit.

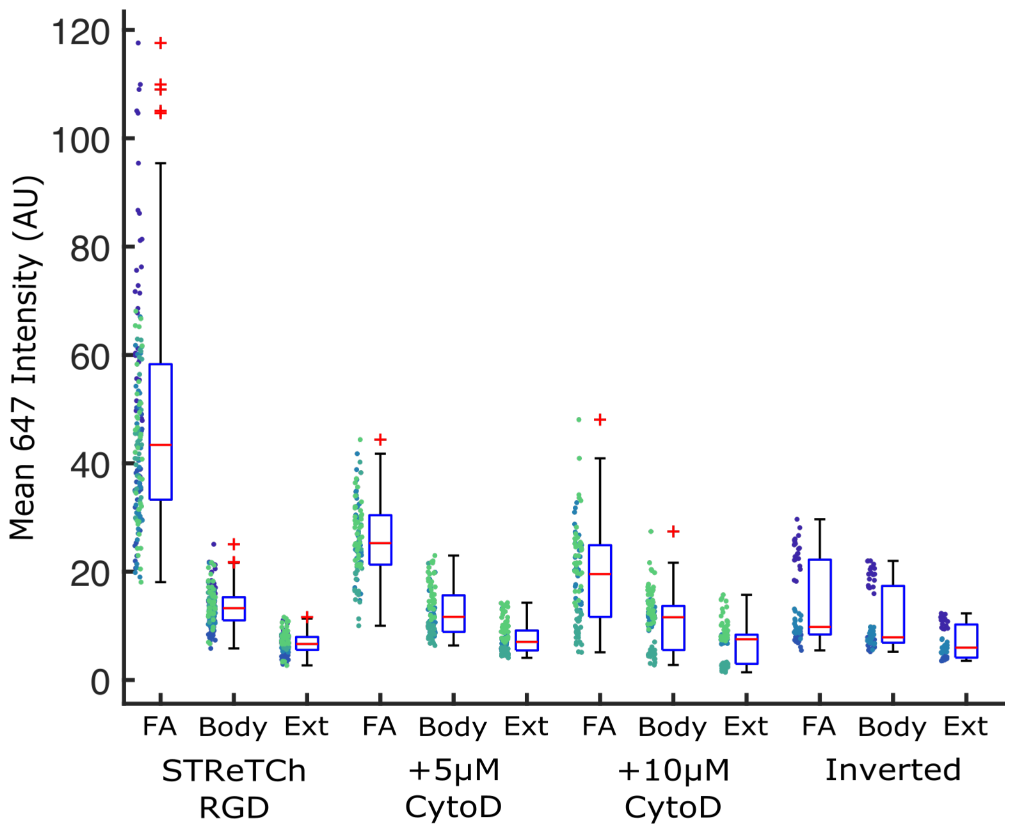


**Supplementary Figure 3. Mean intensities in specific cell regions for extracellular force measurements in Figure 2**

Mean SC-647 intensities in focal adhesions (FAs), outside FAs but underneath cells (Body), and outside cells (Ext) for SC-647 treated HFFs adhering to STReTCh-RGD (with or without Cytochalasin D) or the force-inert inverted control. These data were used to compute the ratios depicted in Fig. 2c and 2d. Red lines indicate medians. Top and bottom of blue boxes indicate 75$^{th}$ and 25$^{th}$ percentiles, respectively, black bars indicate range (excluding outliers), and outliers are plotted as red plus signs. All subsequent boxplots follow this convention. Intensities for individual cells are plotted to the left of each bar, and data from the same experimental trial are indicated with the same color. Apparent bimodality of certain data is due to variation across trials rather than variation among cells within a single trial.

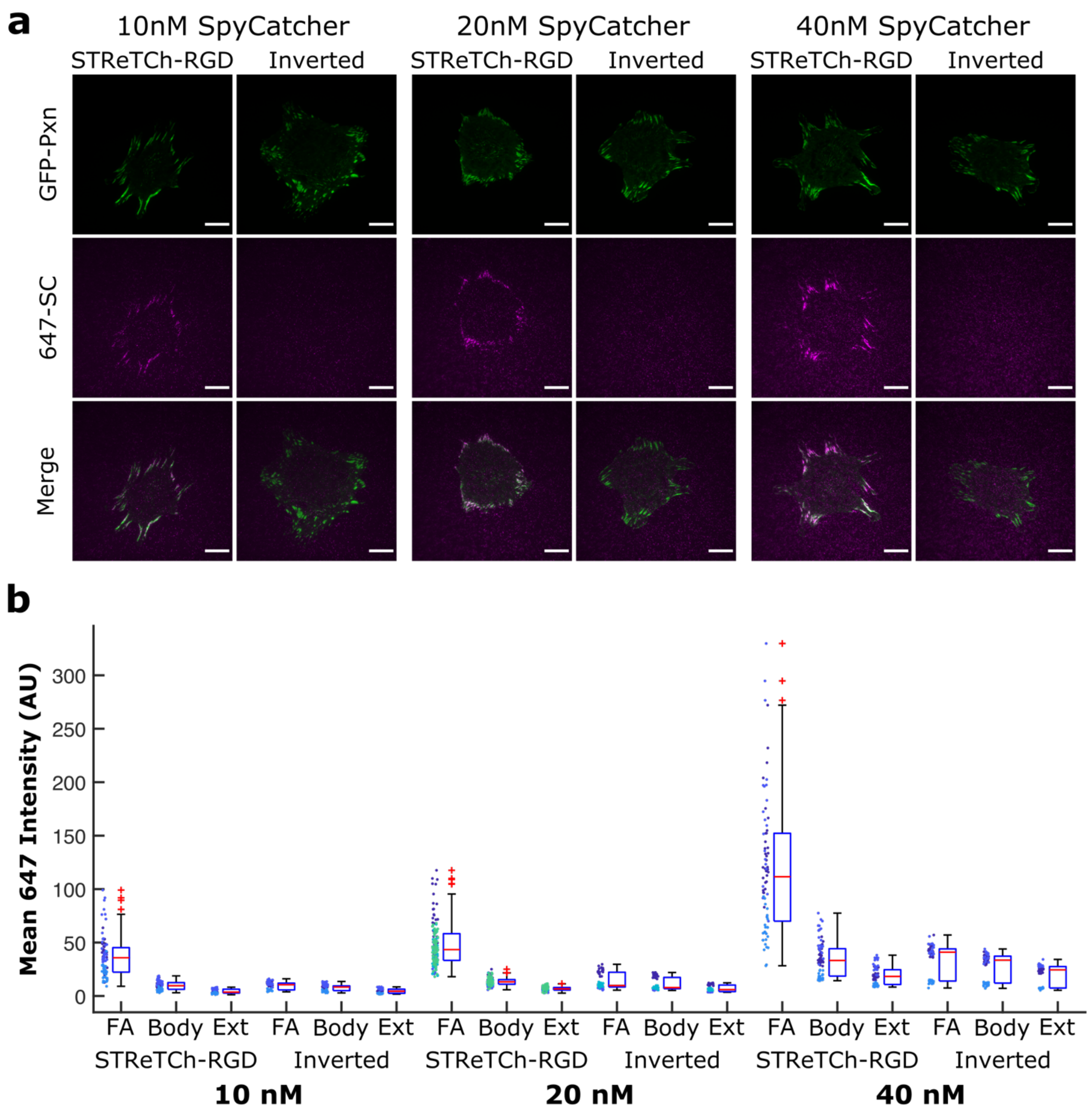


**Supplementary Figure 4. Visualizing force exerted by HFFs on STReTCh-RGD at varying SpyCatcher concentrations.**

**(a)** Representative images of lightly fixed HFFs adhering to STReTCh-RGD or inverted controls when stained with 10, 20, or 40 nM of 647-SpyCatcher for 10 minutes. Scale bars = 20 µm. **(b)** Quantification of mean 647-SC intensity in FAs, outside FAs but beneath cell bodies, and outside of cells for conditions illustrated in **(a)**. Intensities for individual cells are plotted to the left of each bar, and data from the same experimental trial are indicated with the same color. Apparent bimodality of certain data is due to variation across trials rather than variation among cells within a single trial. *N* = 84 cells for 10 nM STReTCh-RGD, 60 for 10 nM inverted, 162 for 20 nM STReTCh-RGD (same dataset as in Fig. 2), 59 for 20 nM inverted (same dataset as in Fig. 2), 78 for 40 nM STReTCh-RGD, and 61 for 40 nM inverted. Data for each condition are pooled from a minimum of 3 independent experiments.

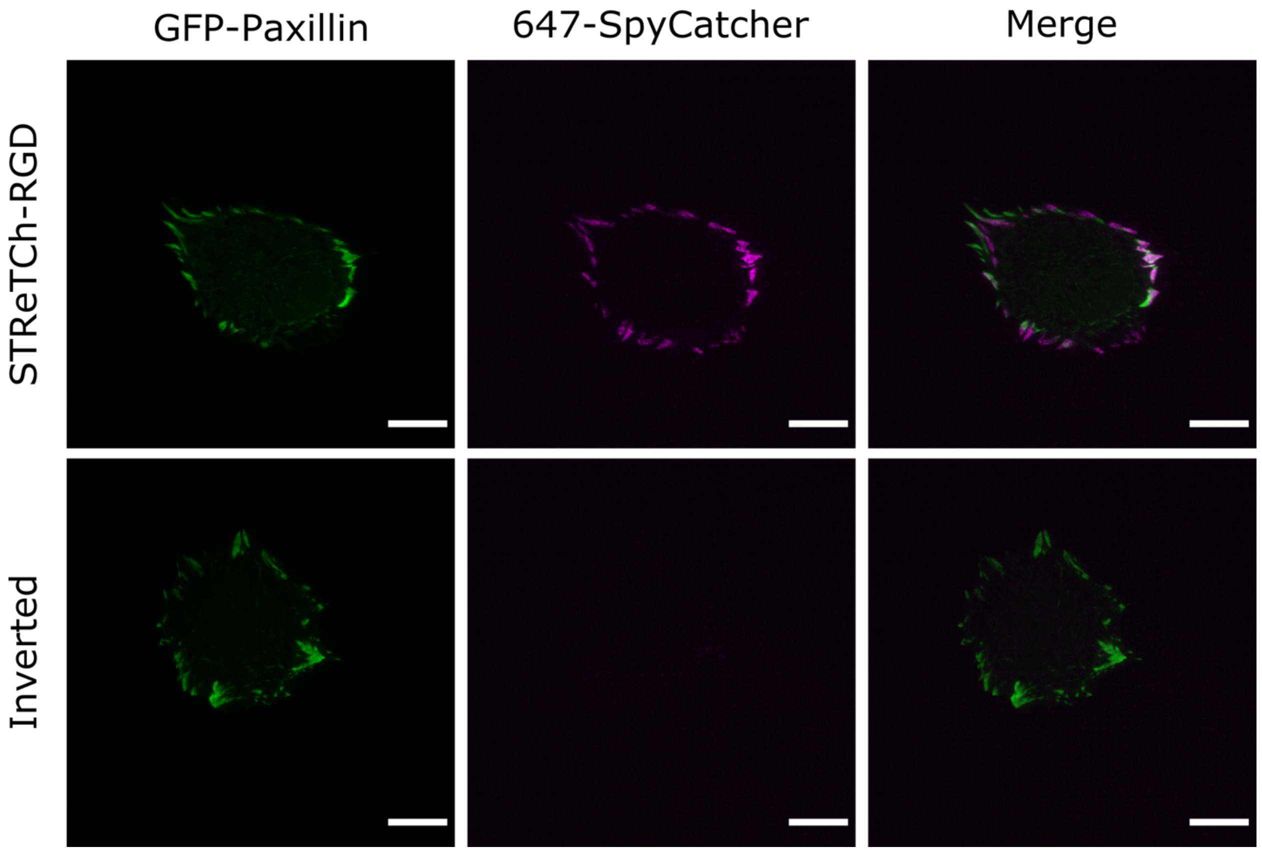


**Supplementary Figure 5. Measurements of extracellular force in living cells**

Representative TIRF microscopy images of live GFP-Paxillin HFFs adhering to STReTCh-RGD (top) and inverted control sensor (bottom) stained with 647-SpyCatcher and imaged live. GFP-Pxn and 647-SC signal on STReTCh-RGD is somewhat spread out due to turnover of FAs during live-cell imaging. Scale bars, 20 μm.

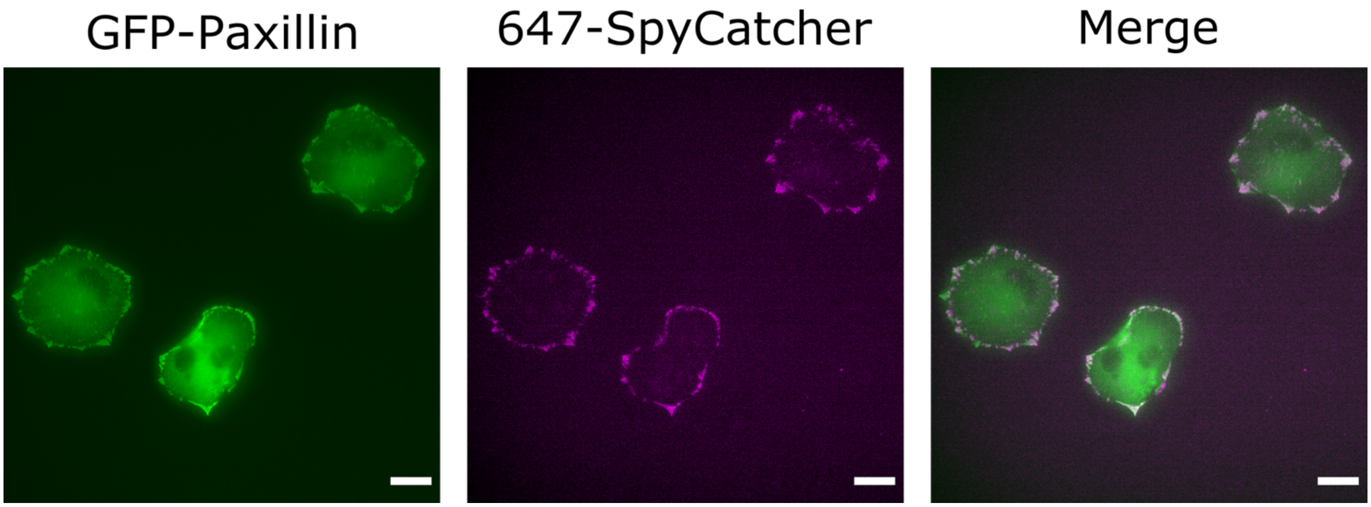


**Supplementary Figure 6. Epifluorescence images of extracellular force sensing for HFFs adhering to STReTCh-RGD**

Representative images of GFP-paxillin (*left*), 647-SpyCatcher (*middle*), and merged (*right*) for GFP-Pxn HFFs adhering to STReTCh-RGD, stained with 647-SC and imaged live using an epifluorescence microscope. Scale bars = 20 μm.

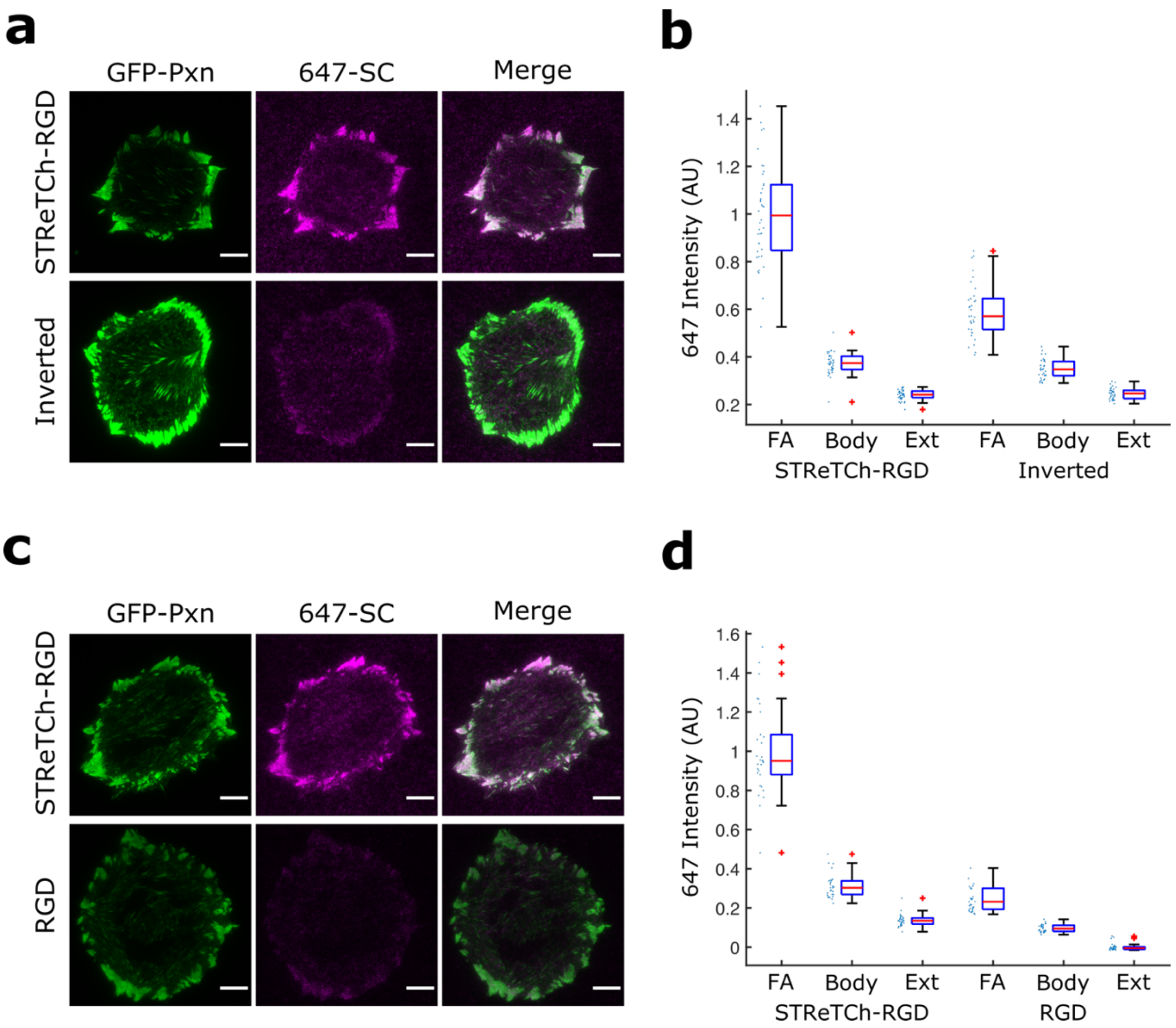


**Supplementary Figure 7. Non-specific background sticking of SpyCatcher to FAs in fixed and permeabilized HFFs**

**(a)** Representative TIRF images of GFP-paxillin and 647-SC in HFFs seeded on STReTCh-RGD or the inverted control sensor, fixed, permeabilized, stained with 647-SC, and then washed with PBS. Scale bars = 10 µm **(b)** Quantification of 647-SC signal intensity in FAs, underneath the cell body but outside of FAs, and outside the cell body for HFFs in (a). *N* = 40 cells for STReTCh-RGD and 34 for inverted. Data are pooled from 2 independent experiments **(c)** Same as (a) but for HFFs adhering to either STReTCh-RGD (*top*) or a flagelliform protein-RGD construct that does not contain SpyTag (bottom)[2-4]. **(d)** Same as (b) but for cells in (c). *N* = 30 for STReTCh-RGD and 31 for RGD. Data are from a single experiment. Intensities are background subtracted using rolling ball background subtraction for (a) and (b) and using the mean intensity outside of cells in the RGD condition for (c) and (d) due to the lower mean intensity in the RGD condition, likely owing to the lack of a reactive SpyTag. These background-subtracted intensities were then normalized to the average FA intensity in HFFs on STReTCh-RGD for each experimental replicate to give the values in (b) and (d).

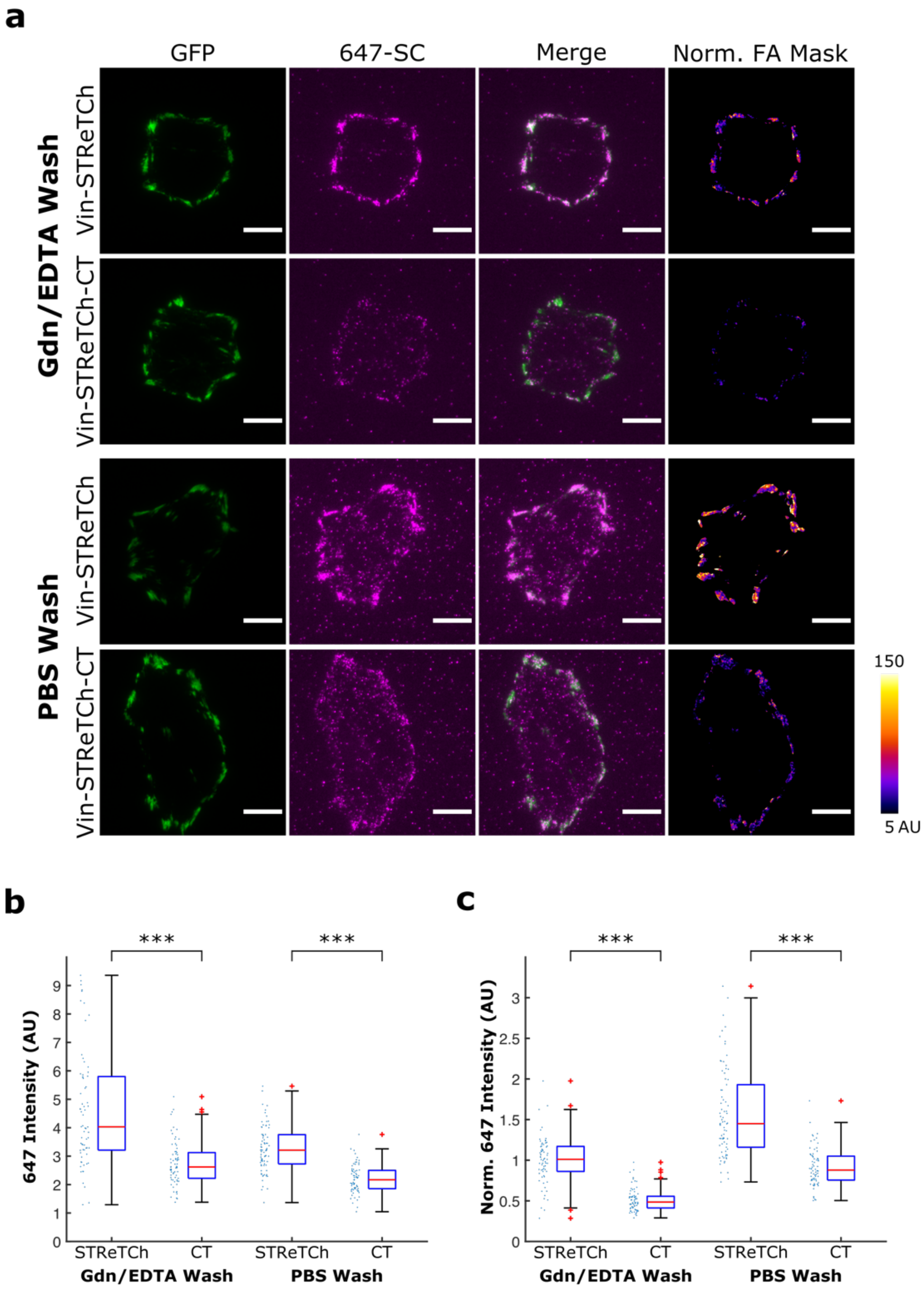


**Supplementary Figure 8. Comparison of force visualization results in Vin-STReTCh and Vin-STReTCh-CT MEFs**

**(a)** Representative images of Vin-STReTCh and Vin-STReTCH-CT MEFs stained with 647-SC and washed with either 3M Gdn-HCl and 2 mM EDTA or with PBS. Scale bars = 10 µm. **(b)** Ratios of 647-SC intensity in FAs and within the cell body but outside of FAs for conditions depicted in (a). **(c)** Quantification of 647-SC intensity normalized by GFP intensity within FAs for conditions depicted in (a). Gdn-HCl/EDTA wash data are reproduced here from Fig. 3. For PBS washes, *N* = 78 cells for STReTCh and 70 for STReTCh-CT and data are pooled from 3 independent experiments. *** denotes $p < 0.001$ by two-tailed Mann-Whitney test.

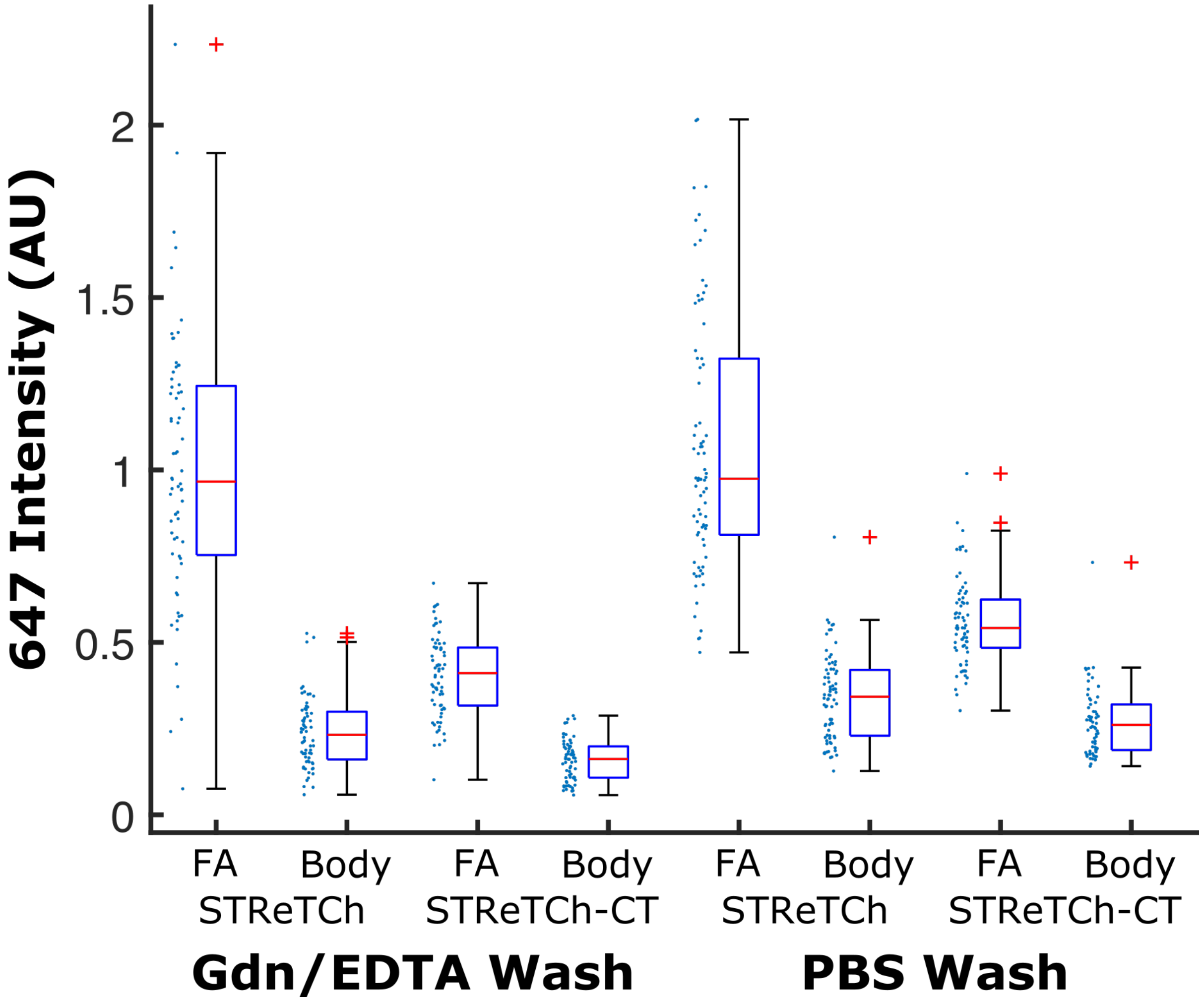


**Supplementary Figure 9. Mean intensities in specific cell regions for intracellular force measurements across vinculin in MEFs.**

Mean 647-SC intensities above background (cell exterior) in FAs and outside FAs but underneath cells (Body) for Vin-STReTCh and Vin-STRetCh-CT MEFs washed with either 3 M Gdn-HCl and 2 mM EDTA or PBS. These data were used to compute the ratios depicted in Fig. 3c,d and Supp. Fig. 8b, c. For each of 3 independent experiments, mean 647-SC intensity values for each condition were normalized to the average of mean 647-SC intensity values within FAs for that specific experiment. These normalized values are then pooled to give the values depicted in this figure.

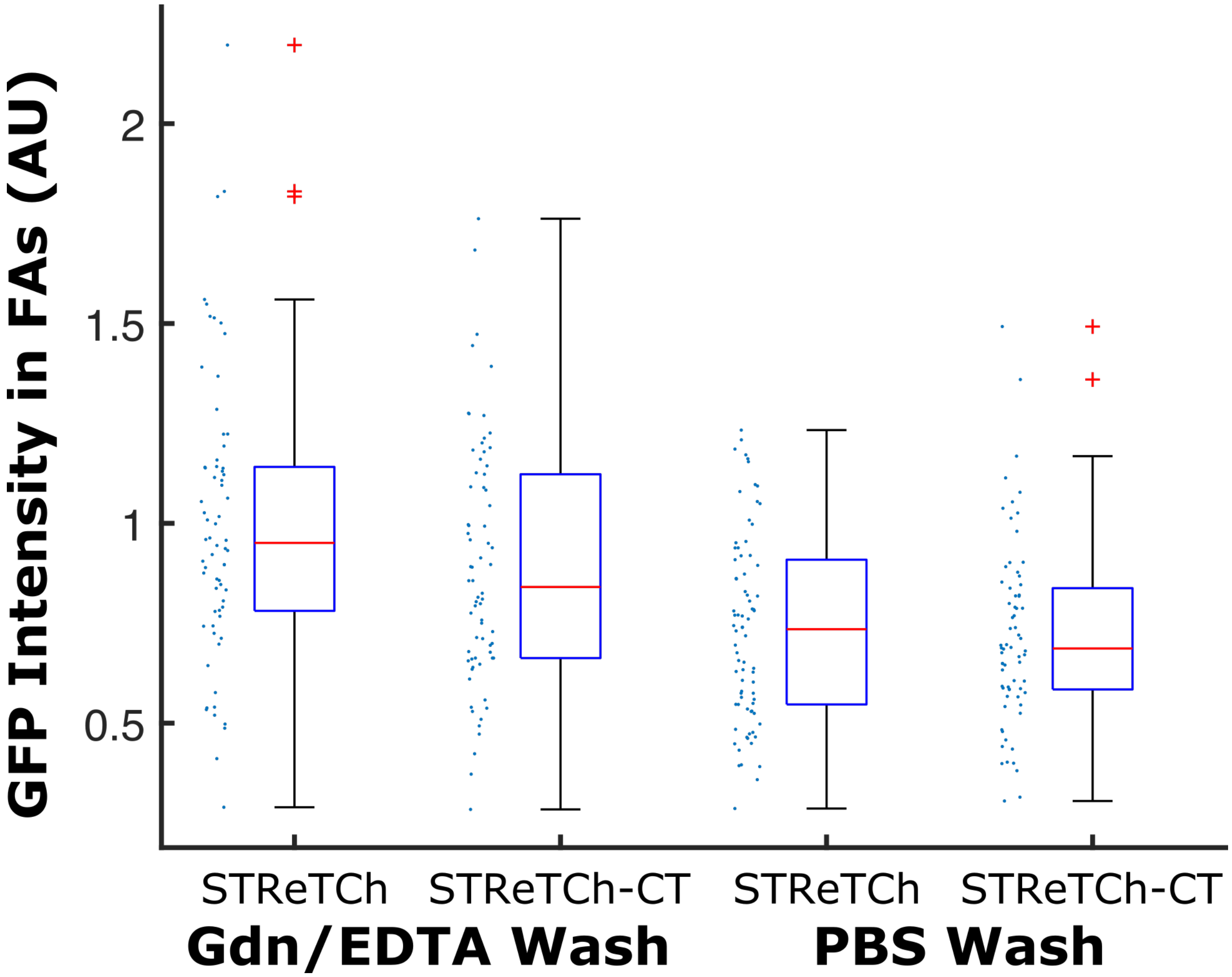


**Supplementary Figure 10. GFP Expression at FAs for Vin-STReTCh and Vin-STReTCH-CT MEFs**

Vin-STReTCh and Vin-STReTCh-CT MEFs analyzed in this study show comparable levels of construct expression and localization at FAs, as measured by GFP intensity at FAs. $p = 0.062$ for STReTCh vs STReTCh-CT in Gdn/EDTA wash (Fig. 3, Supp. Fig. 8), and $p = 0.62$ for PBS wash (Supp. Fig. 8) by two-tailed Mann-Whitney test.

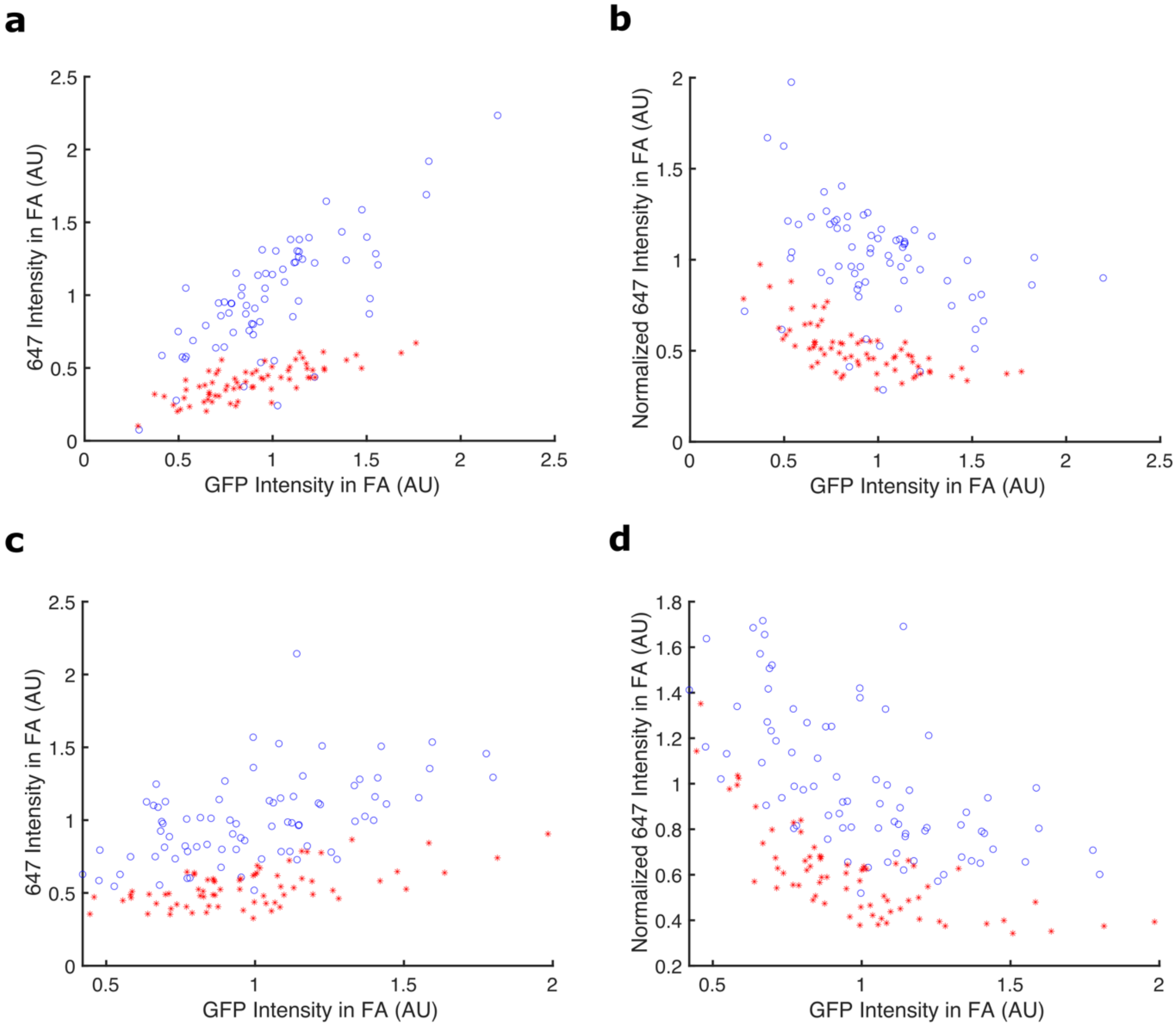


**Supplementary Figure 11. 647 intensity and normalized 647 intensity within FAs in Vin-STReTCh and Vin-STReTCh-CT MEFs plotted against GFP intensity within FAs**

**(a)** 647-SC intensity in FAs for Vin-STReTCh (*blue*) and Vin-STReTCh-CT (*red*) MEFs (Fig. 3, Supp. Fig. 8) plotted against average FA GFP intensity. Each point represents one cell and cells were washed with 3 M Gdn-HCl and 2 mM EDTA. **(b)** 647-SC intensity in FAs normalized by FA GFP intensity for Vin-STReTCh (*blue*) and Vin-STReTCh-CT (*red*) MEFs (Fig. 3, Supp. Fig. 8) plotted against average FA GFP intensity. Each point represents one cell and cells were washed with 3 M Gdn-HCl and 2 mM EDTA **(c)** Same as (a) but for MEFs washed with PBS (Supp. Fig. 8). **(d)** Same as (b) but for MEFs washed with PBS (Supp. Fig. 8).

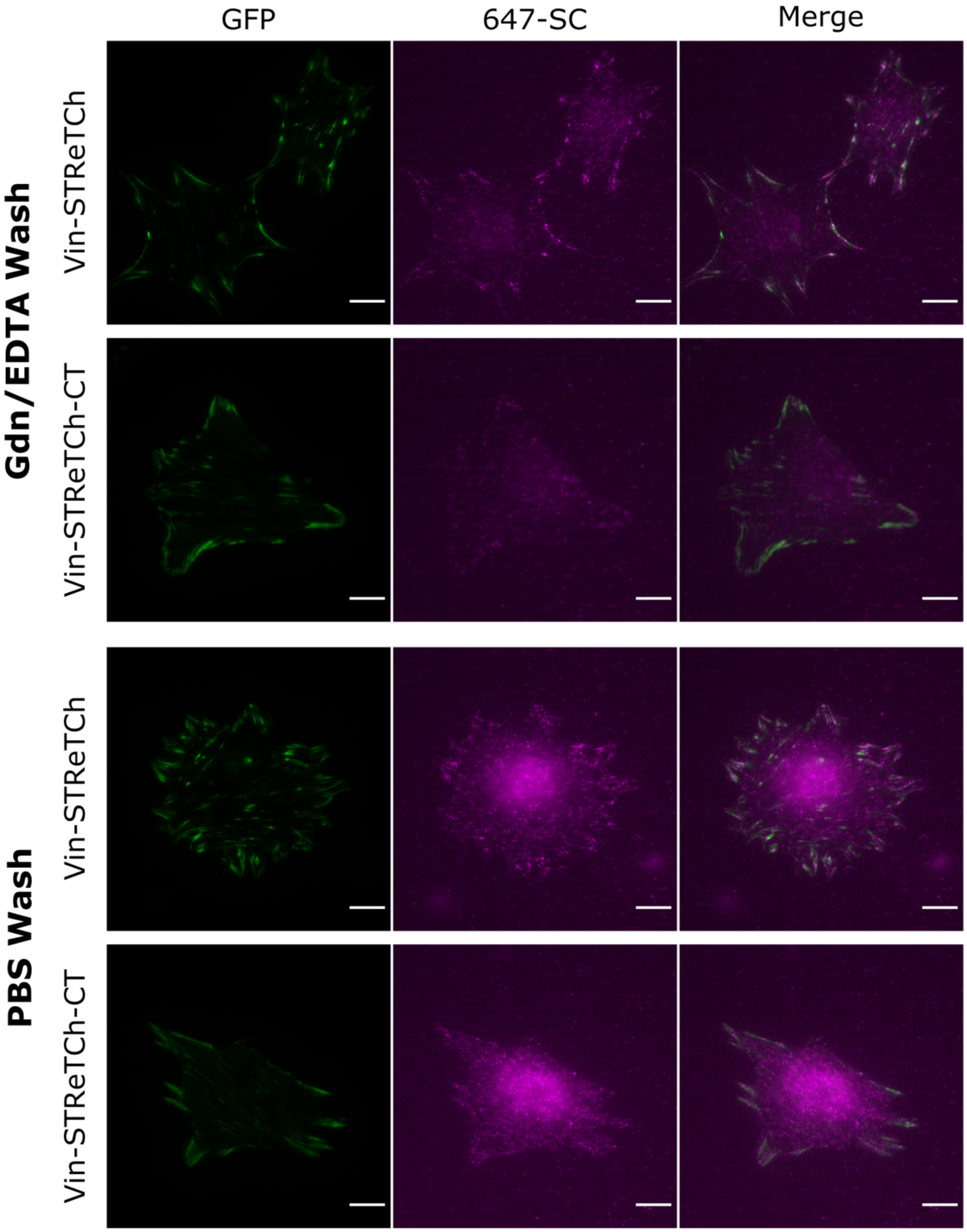


**Supplementary Figure 12. Images of Vin-STReTCh and Vin-STReTCh-CT MEFs stained with 647-SC and imaged using epifluorescence illumination**

Representative images of Vin-STReTCh and Vin-STReTCh-CT MEFs on fibronectin, fixed, permeabilized, and treated with 5 nM 647-SC for 10 minutes, then washed with either 2 mM EDTA and 3 M Gdn-HCl (top) or with PBS (bottom). Images were acquired by adjusting the TIRF angle from the TIRF microscopy setup (see Materials and Methods) to provide direct epifluorescence illumination to the sample. Scale bar = 10 μm.

**Supplementary Table 1. Maximum likelihood parameter estimates for Bell-Evans fit to magnetic tweezers data**

| | Estimate | Lower 95% CI | Upper 95% CI |
|---|---|---|---|
| $k_u^0$ (s$^{-1}$) | 2.92x10$^{-3}$ | 4.88x10$^{-4}$ | 1.44x10$^{-2}$ |
| $\frac{\Delta x_u}{k_B T}$ (pN$^{-1}$) | 1.12x10$^{0}$ | -3.25x10$^{-1}$ | 2.59x10$^{0}$ |
| $k_f^0$ (s$^{-1}$) | 6.59x10$^{-2}$ | 1.00x10$^{-2}$ | 3.91x10$^{-1}$ |
| $\frac{\Delta x_f}{k_B T}$ (pN$^{-1}$) | -1.89x10$^{0}$ | -3.52x10$^{0}$ | -3.39x10$^{-1}$ |
| $F_{50\%}$ (pN) | 1.03 | 0.19 | 3.25 |
